# Energy-based bond graph models of glucose transport with SLC transporters

**DOI:** 10.1101/2024.06.26.600892

**Authors:** Peter J. Hunter, Weiwei Ai, David P. Nickerson

## Abstract

The SLC (<u>s</u>olute <u>c</u>arrier) superfamily mediates the passive transport of small molecules across apical and basolateral cell membranes in nearly all tissues. In this paper we employ bond graph approaches to develop models of SLC transporters that conserve mass, charge and energy, respectively, and which can be parameterised for a specific cell and tissue type for which the experimental kinetic data is available. We show how analytic expressions that preserve thermodynamic consistency can be derived for a representative four- or six-state model, given reasonable assumptions associated with steady-state flux conditions. We present details on fitting parameters for *SLC2A2* (a GLUT transporter) and *SLC5A1* (an SGLT transporter) to experimental data and show how well the steady-state flux expressions match the full kinetic analysis. Since the bond graph approach will not be familiar to many readers, we provide a detailed description of the approach and illustrate its application to a number of familiar biophysical processes.

**SIGNIFICANCE:** Physiological systems typically involve coupled mechanical, electrical and chemical processes, with energy acting as a universal currency across these domains. We propose a new visual representation for all components of these processes using bond graphs. Bringing all physical processes under one consistent framework greatly simplifies the task of understanding multiscale physiological processes. This energy-based framework, which is the 0D version of a more general 3D port-Hamiltonian theory, can be used to model all lumped parameter physiological processes. A small number of bond graph templates can be used to model all members of the large SLC transporter family, and reduced thermodynamically consistent steady-state flux models provide a useful simplification for many situations. Glucose transport is chosen here to illustrate the bond graph approach because it represents the first step in cell metabolic processes, where energy conservation needs to be a fundamental characteristic of quantitative models. Our future work on cell metabolism will build on the foundation established here.

## INTRODUCTION

The solute carrier (SLC) superfamily currently consists of proteins encoded by more than 400 mammalian genes that mediate the transport of small molecules across cell and organelle membranes in human tissues [1]. ATP-dependent pumps, ATP-binding cassette transporters, aquaporins and ion channels belong to separate families of transport proteins, which together comprise at least 5% of the protein-coding genome. The SLC superfamily is currently categorised into 62 gene families, labelled *SLC1* to *SLC62* (www.bioparadigms.org). Two or more families may deal with transport of the same ligand (e.g. glucose) but each family deals with a specific type of transport mechanism. For example, transmembrane glucose transport across a number of cell types (endothelial, epithelial, neuronal, etc) is enabled by two families of protein from the SLC superfamily: the *SLC2* family and the *SLC5* family. *SLC2A2* (protein name GLUT2) and its variants within that family use the extracellular to intracellular glucose concentration gradient to drive transmembrane transport in a process called ‘facilitated diffusion’. No other ligands are involved. On the other hand, *SLC5A1* (SGLT1) and its variants use the sodium gradient to drive glucose into the cell, typically when the transmembrane glucose gradient is insufficient to provide the required flux of glucose.

Bond graphs provide a useful level of abstraction for modelling protein function for a wide range of physiological processes, such as metabolic reactions, membrane transporters, ion channels, myofilament mechanics, receptors and signalling, etc. In this paper we develop a small number of generic bond graph templates for the SLC superfamily that conserve mass, charge and energy, respectively, and which can be parameterised for a specific cell and tissue type for which the experimental kinetic data is available. We show how analytic expressions can be derived for a representative six-state model, given reasonable assumptions associated with steady-state flux conditions, while always preserving thermodynamic consistency. We present details on fitting parameters for *SLC2A2* and *SLC5A1* to experimental data and show how well the steady-state flux expressions match the full kinetic analysis.

Before developing the SLC family templates, we first discuss some fundamental physical concepts and their units, describe why bond graphs provide an appropriate framework for capturing the physical conservation laws associated with biological processes at the protein level, and present some examples that illustrate how bond graphs deal with energy exchange. Bringing all physical processes under one consistent framework greatly simplifies the task of understanding physiological processes, which almost always involve energy exchange between mechanics, electromagnetics and chemistry.

## METHODS

### Units, conservation laws and bond graphs

We start by discussing the key units for physiology. Only six fundamental units (Joules, entropy, seconds, meters, Coulombs, and moles) are needed for all biophysical mechanisms, with energy gradients (with respect to meters, Coulombs and moles) providing the driving ‘force’ or potential for displacement from equilibrium for each of the three physical processes that underpin physiology: *mechanical* (J.m^-1^ or J.m^-3^), *electromagnetic* (J.C^-1^), and *chemical* (J.mol^-1^). Note that the Coulomb (C) effectively counts electrons and the mole (mol) counts atoms. Very occasionally it is useful to include a seventh unit, the Candela (Cd), which counts photons (for example in models of photoreceptors that respond to individual photons) but generally a photon (which has an energy *hv*, or Planck’s constant *h* times the frequency *v* of the electromagnetic field) is included via its energy flux. Energy, measured in Joules (J), can be transmitted, stored, or converted between these three types, and almost every physiological process uses all of them. The closely related concept of entropy is a measure of displacement from equilibrium and energy dispersion (or equivalently the possible states of a system). Enthalpy *H* is defined as the sum of internal energy *U* (associated with vibrational, rotational and electronic states of the molecules) and the product *pV* of thermodynamic pressure *p* (an energy density) and volume *V* (*pV* = *nRT* for an ideal gas), but is also the sum of the Gibbs free energy *G* (available to do work) and the *TS* term representing the essential loss of high entropy energy (heat) to the environment:

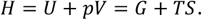

Total energy is conserved (but not *G*). For example, a thermally stable planet earth must receive and radiate energy at the same rate, but for every 1 high energy (∼500nm wavelength) photon that the earth receives as highly directed (low entropy) sunlight, it radiates about 20 times as many lower energy (∼10,000nm wavelength) photons as (high entropy) heat. This conversion of conserved energy from low to high entropy form defines the direction of time and is of course the basis for life. It is the *energy gradient* with respect to meters, Coulombs or moles that is the ‘*potential’* (i.e. the mechanical force, electrical potential or chemical potential) that drives the flow or flux of a mechanical, electrical or chemical quantity. Note that temperature, measured in degrees Kelvin, is the gradient of energy with respect to entropy, and hence is the thermal potential driving heat flow (the flow of entropy). It is convenient to define the unit of entropy as Joules per degree Kelvin, since it is impractical to count the number of possible states in a thermally energetic system.

Energy storage is either *mechanical* (statically in a spring or dynamically in the inertia of a mass), *electromagnetic* (statically in a capacitor or dynamically with the inductance of a changing magnetic field), or *chemical* (statically as a solute in a solution or dynamically as thermal energy). Note that processes at the macroscale of physiology are linked with processes at the atomic level through a small number of physical constants such as Faraday’s constant (9.6485×10^4^ C.mol^-1^; the charge in Coulombs of a mole of single charge ions) and the gas constant (the energy in Joules per degree Kelvin of a mole of atoms), both of which use Avogadro’s number (6.02214 x10^23^ mol^-1^) to bridge the enormous scale from atoms to cells and tissues.

We let *q* (in units of m, m^3^, C, or mol) represent the quantity whose flow *v* (in units m.s^-1^, m^3^.s^-1^, C.s^-1^, or mol.s^-1^) is driven by a potential *u* (in units J.m^-1^, J.m^-3^, J.C^-1^, or J.mol^-1^). We use two forms of mechanical flux (in m.s^-1^ and m^3^.s^-1^) with potentials in J.m^-1^ (Newton) and J.m^-3^ (Pascal), in order to deal with both solid mechanics and fluid mechanics. Note that for the most part there is no need to use any derived units (such as the Newton or Pascal). Using only J, K, s, m, C and mol helps reinforce the relationships between these units that underpin both the conservation laws of physics and the constitutive laws that represent material properties.

There are two distinct types of equation needed for characterising physical systems (note that we provide explicit examples of electromechanical and biochemical processes in the next section). The first type is a *physical conservation law* (conservation of mass, charge or energy, respectively), which generates equations that involve only *q* or *v* (mass or charge conservation), or only *u* (energy conservation). The second type is a *constitutive equation* that expresses experimentally derived material properties and is an equation that links *q* or 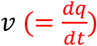 with *u*. These material properties relate to either (i) *energy storage*, which is a relationship between *u* and *q* for static storage, or *u* and 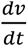 for dynamic storage, (ii) *energy dissipation* (a mechanical damper, an electrical resistance, or a chemical reaction), which is a relationship between *u* and *v*, or (iii) lossless *energy conversion* between mechanical, electromagnetic and chemical energy. Physical conservation laws are universally true, whereas the constitutive equations are approximations fitted to experimental data (and hide the physics behind that material behaviour). The wide variety of ad-hoc descriptions of physical processes often presented in physiology textbooks, such as Fick’s law of diffusion, Fourier’s law of heat conduction, or osmotically driven flow, etc, express a combination of these fundamentally different types of equation in a way that conflates the laws of physics with experimentally derived material properties.

The conservation laws that govern all physical processes, and the empirical constitutive equations, can both be expressed with the above quantities *q*, flows *v* and potentials *u* in a very simple, elegant and unifying manner by using a technique called *bond graphs*, pioneered for electromechanics by Henry Paynter at MIT [2]. In another fundamental and far-sighted contribution, Oster, Perelson and Katchalsky [3, 4] brought network thermodynamics within the same framework so that there is now a single unifying energy-based framework for all of physics at the spatial scales relevant to physiological mechanisms. The physiological application of bond graphs began with a series of papers by Gawthrop and Crampin [5, 6], who demonstrated the importance of energy conservation in modelling physiological mechanisms.

The key concept is this: the product of potential *u* and flow *v* is power *u*. *v* in units of J.s^-1^. Paths for the transmission of power, called *bonds*, are shown by the directed arrows in Figure 1a (the arrow defines the direction of positive power flow). Each bond with subscript *i* carries a flow *v*_*i*_ at potential *u*_*i*_. At the junction of bonds, conservation of power requires:

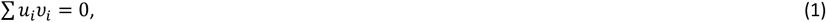

where summation is carried out over all bonds connected to that junction (5 in this example). Now consider the case shown in Figure 1b (called a *0-node*) where all bonds have the same potential *u*_1_ = *u*_2_ = *u*_3_ = *u*_4_ = *u*_5_ = *u*, in which case *u* comes outside the summation and, for non-zero *u*, the conservation of power becomes conservation of flow:

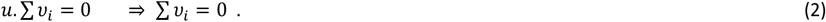

If *u*_*i*_ represents the flow of a volume of fluid, and if the density of the fluid is constant (which it is for water in physiological systems), equation 2 represents conservation of mass. If *u*_*i*_ represents the flow of charge, equation 2 represents conservation of charge, etc.

**Figure 1.**
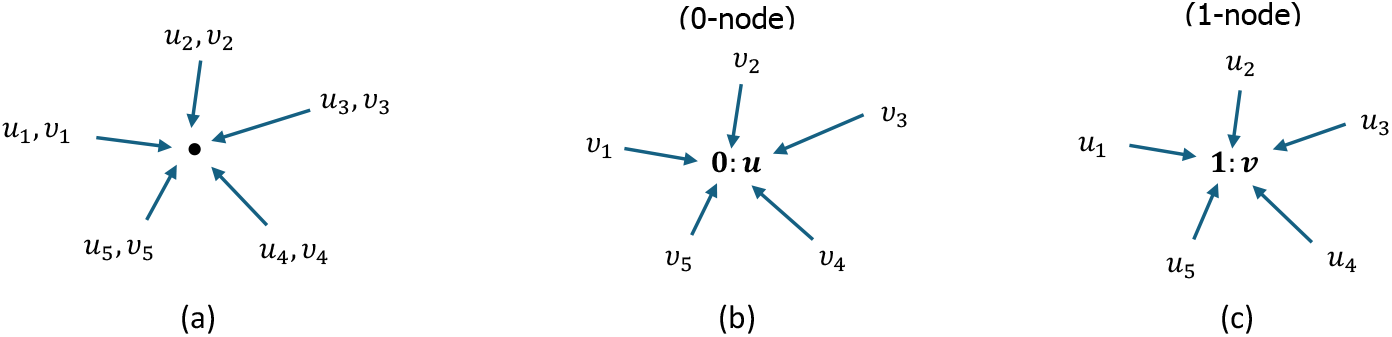
Bond graphs representing power flow, each defined with values for flow *υ*_*i*_ and potential *u*_*i*_: (a) a junction of 5 flow paths such that ∑ *u*_*i*_*υ*_*i*_ = 0, (b) a junction where the impinging bonds have the same potential *u* (called a 0-node) and hence ∑ *υ*_*ii*_ = 0, and (c) a junction where the impinging bonds have the same flow *v* (called a 1-node) and hence ∑ *u*_*i*_ = 0. Therefore 0-nodes are junctions on a bond graph where quantities are conserved (mass, charge, etc) and 1-nodes ensure that energy is conserved.

Alternatively, consider the case shown in Figure 1c (called a *1-node*) where all bonds have the same flow *v*_1_ = *v*_2_ = *v*_3_ = *v*_4_ = *v*_5_ = *v*, in which case *v* comes outside the summation of equation 1 which, for non-zero *v*, becomes conservation of energy:

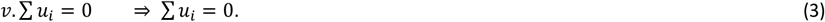

In summary, power transmission is modelled here via bond graphs that converge on power-conserving junctions of just two types: 0-node junctions with a common potential are points at which flows sum to zero so that a quantity (mass, charge, etc) is conserved, and 1-node junctions with a common flow are points at which the potentials sum to zero and energy is therefore conserved. The variables *u* and *v*, whose product is power, are called power *co-variables* or *conjugate variables*. Note that bond graphs describe the topology of the system (i.e., how the components are connected) as well as the equations that capture the conservation laws of nature. Power is always conserved but the ability to solve the bond graph system (and the solution itself) is determined by the boundary conditions.

Now consider the static storage of energy. Since power (*u*. *v*) is the rate of change of energy,

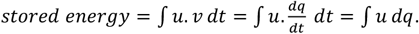

For linear storage devices, *u* = *C*^−1^*q*, where compliance *C* is an empirically determined constant,

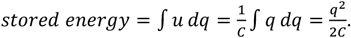

The relationship *u* = *f* (*q*) is the *constitutive relation* for that material. E.g., for a linear mechanical spring, the mechanical potential *u* (force) is proportional to the displacement *q* of the spring, and *C*^−1^ is the spring stiffness. Exactly the same linear expression holds for an electrical capacitor: *q* is charge, *u* is the electrostatic potential (voltage) across the capacitor and *C* is now the capacitance. Biochemical storage depends on the solubility of the solute in the solvent.

Next we consider the mechanisms involving dissipation of energy (and hence the production of heat). In most cases the rate of energy dissipation is just the product of the flow through the dissipator and the change in potential across it (e.g., a viscous damper in mechanics, a resistor in an electrical circuit, or a thermal resistance in heat flow). For most situations the drop in potential is assumed to be linearly proportional to flow (Δ*u* = *Rv*), giving

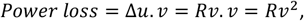

where the experimentally determined constitutive parameter *R* is termed the mechanical viscosity, electrical resistance, or thermal resistance.

The dissipative mechanism in biochemistry is a chemical reaction and for this case it has a special form which is both nonlinear and depends explicitly on the forward and reverse affinities (*A*^*f*^ and *A*^*r*^), representing the sum of chemical potentials for the reactants (*u*_1_) and products (*u*_2_), with the Marcelin-de Donder formula [5]:

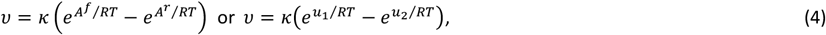

where the specific reaction rate *k* has units of mol.s^-1^. Note that the flow is not determined by just the difference in potentials as in all the other systems but rather uses an explicit dependence on each (and hence is called a ‘two-port’ device). Reactions will only proceed if *u*_1_ > *u*_2_, or equivalently if the Gibbs free energy Δ*G* = *u*_2_ − *u*_1_ < 0. However, the power that is emitted as heat in a biochemical reaction is still the same product *v*. Δ*u* (= −*v*. Δ*G*) as it is in an electrical resistor or mechanical damper.

A chemical reaction requires the exchange or sharing of electrons (it is a sub-atomic process) and since all members of the chemical species involved in that reaction are available in a well-mixed compartment, the chemical potential energy is normalised by the total number of moles in that compartment (i.e. is an intensive property). A diffusion process, on the other hand, is entropically driven and entropy is an extensive property. Unlike engineering processes, where the heat from an electrical resistor or a mechanical damper is generally lost to the environment, the heat output *v*. Δ*u* from a biochemical reaction in a physiological system is used for temperature regulation.

The process of transforming power without loss from one physical process to another (as in a voice coil or ‘loudspeaker’, where electrical power is transformed to mechanical power) is illustrated in Figure 2a. The electrical power co-variables associated with the first bond are 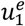 and 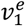 and the mechanical power co-variables associated with the second bond are 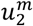 and 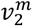. We define an empirical constitutive relation in which the output mechanical potential 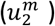 is proportional to the input electrical current flow 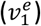:

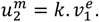

Since power is conserved, 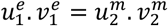, and a rearrangement gives

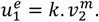

The first of these equations is the Lorenz force on a voice coil and the second is Faraday’s ‘back-EMF’ (electromotive force) induced by movement of the coil. The empirical parameter *k* is associated with the ‘Gyrator’ term GY defined at the junction in Figure 2a. The conversion of electrical energy in J.C^-1^ to chemical energy in J.mol^-1^ is another example of the need for the gyrator term *k* (see later).

**Figure 2.**
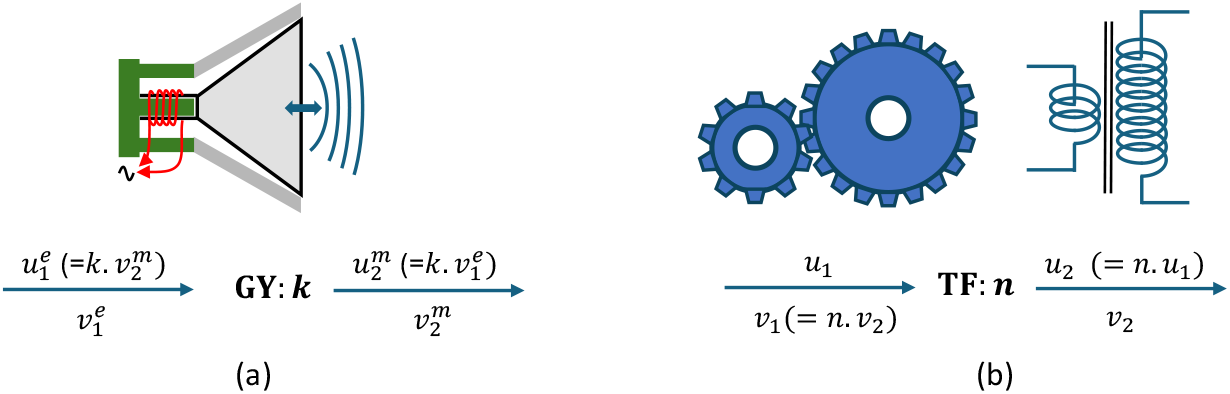
Transforming energy (without loss) (a) from one form to another (electrical to mechanical here), and (b) at different ratios of flow to potential (e.g., between two gear wheels or two transformer windings). Note that the coil in (a) is attached to the moving cone of the speaker.

Another form of transformation in which potential is traded for flow in a way that conserves power is shown in Figure 2b. The ‘transforming factor’ (TF) is associated with a dimensionless parameter *n* representing the n-fold increase in output potential and corresponding n-fold decrease in output flow. Examples are a mechanical gear wheel, an electrical transformer, and a mechanical lever.

A bond graph diagram contains all the information needed to create the model and is a very convenient way to visualise the energy transmission, energy storage, and energy conversion (including dissipation to heat) occurring in the system being modelled.

### Simple bond graph examples

Before addressing the SLC family of transporters, we show how bond graphs are used to create models based on physical conservation laws for three simple examples, one of a coupled electromechanical actuator (a voice coil), one of a voltage-sensitive and mechano-sensitive gated ion channel, and one of an enzyme-catalysed reaction. These examples are used both to introduce the graphical nature of bond graphs, including their particular symbols, and to demonstrate how straightforward it is to generate models that obey the three conservation laws of physics, particularly where these models involve the exchange of energy between the three different physical energy storage mechanisms.

#### Example 1: An electromechanical system

The classic example of a coupled electromechanical system is an electrical circuit driving a voice coil (such as a loudspeaker), as shown in Figure 3a. In this case we specify the input voltage 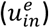 which produces an electrical current 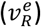 flowing through a resistance R_1_, an inductance L_1_, and the voice coil (length *l*) to produce a time-varying magnetic field of flux density *B* (Js.C^-1^.m^-2^) which generates a (‘Lorentz’) force 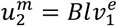 and hence displacement 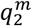 (velocity 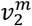). The mechanical system also has a spring of compliance C_2_, a viscous damper R_2_, and an inertia L_2_. The bond graph representation of this system is shown in Figure 3b.

**Figure 3.**
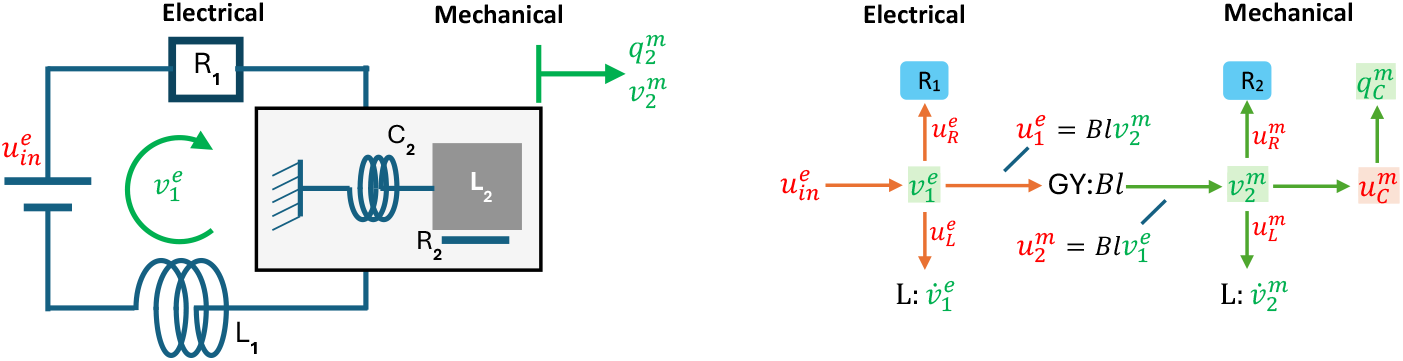
(a) A coupled electromechanical system, and (b) its bond graph representation. 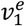 is an electrical current and 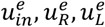 and 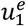 are electrical potentials (voltages). 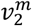 is a mechanical velocity (displacement 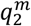) and 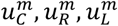 and 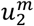 are mechanical potentials (forces).

Note that the coupling between the electrical and mechanical components requires a ‘gyrator’ (GY). The voltage (called ‘back-EMF’) induced within the voice coil is 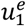 and the expression 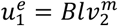 is required to ensure lossless power transfer since the power from the electrical side 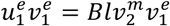 must match the power 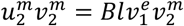 on the mechanical side. Faraday’s law of induction appears as a logical consequence of the Lorentz force. *B* is the magnetic field strength in units of Js.C^-1^.m^-2^.

The balance equations and constitutive laws for this system are:

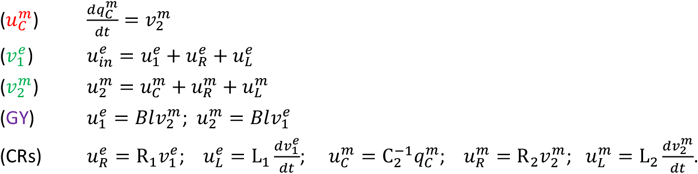

Note that we identify each type of equation using the colours red, green and purple, respectively, for 0:node mass or charge conservation, 1:node energy conservation, and energy conversion. The final equations needed to link the potentials *u* with their appropriate kinematic quantities *q, v* are the constitutive relations (CRs) that capture the material properties of the system components.

R_1_ and L_1_ are the resistance and inductance in the electrical circuit. R_2_, L_2_ and C_2_ are the damping resistance, inertia and spring compliance in the mechanical system. In this bond graph formulation, the Lorentz force and the back EMF from Faraday’s law of induction are just two ways of viewing the same power preserving (GY) mechanism.

With 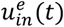 specified as an input condition, these 10 equations can be solved for 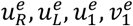 and 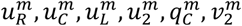.

#### Example 2: A voltage-sensitive and mechano-sensitive gated ion channel

In this example, we consider various physical influences on the movement of charged sodium ions Na^+^ through a membrane ion channel: (i) the chemical potentials associated with different numbers of ions on each side of the membrane; (ii) the effect on a charged ion moving through the electric field associated with the membrane channel; (iii) the effect of membrane stretch (represented by a one-dimensional strain term) on membrane permeability; and (iv) the gating process that controls ion permeation. In general, this gating process is itself voltage-dependent and often subject to ligand binding, but we ignore those factors as the goal here is just to demonstrate the way that a bond graph approach is used to develop an ion channel model that obeys physical conservation laws.

We define two well-mixed (homogeneous) compartments on either side of a semi-permeable channel in the cell membrane, with 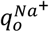 and 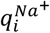 extracellular representing the number of moles of Na^+^ in the space (‘o’=outside) and in the cytosol of the cell (‘i’=inside).

Since biological systems are usually assumed to be at constant temperature and pressure, Gibbs free energy is the relevant chemical potential in these systems. For a dilute system the chemical potential is given (using the extracellular compartment ‘o’ as an example) by the Boltzmann thermodynamic relation

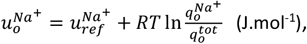

where 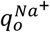 is the number of moles of Na^+^ and 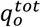 is the total number of moles of all components of the mixture in that compartment [7]. 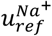 is the (reference) potential when 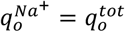.

More compactly,

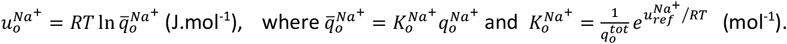

This is the constitutive law for biochemical energy storage, and 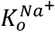 (mol^-1^) is an experimentally determined thermodynamic material parameter. Using the non-dimensional term 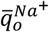 simplifies the subsequent analysis.

Similarly,

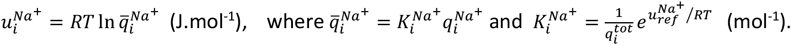

Figure 4 shows the bond graph representation of the diffusive solute flux through the membrane and the flux of electrical charge 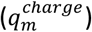 from its capacitive storage in the membrane, with the biochemical equivalent of the membrane potential 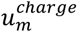 (J.C^-1^) being 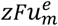 (J.mol^-1^), where *F* is the Faraday constant and *z* is the valence (*z* = 1 for Na^+^).

**Figure 4.**
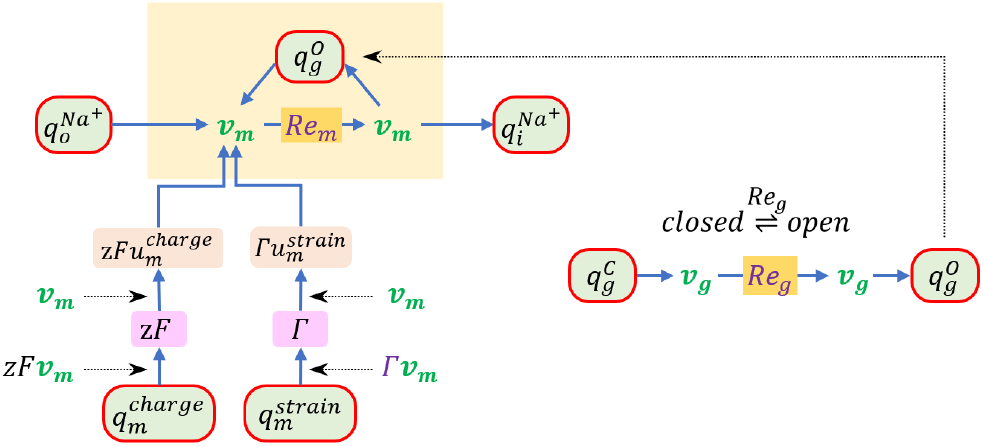
The bond graph representation of the concentration driven flow *v*_*m*_ of electrically charged *K*^+^ ions through a channel in a membrane (inside the yellow block above) across which there is a potential difference 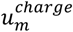 associated with charge storage 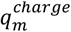. The membrane is also subject to mechanical strain 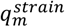. The ion channel is gated by a variable that transitions in reaction *Re*_*g*_ between a closed state 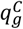 and an open state 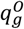. The subscripts are ‘o’ and ‘i’ identify the outside and inside of the cell, respectively, while ‘m’ and ‘g’ refer to the cell membrane and the ion channel gate.

Note that Figure 4 introduces a new symbol with a green background and a red border. The green background indicates that this represents a storage term (see 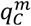 in Figure 3), which determines its potential 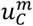, and the red border indicates that this is also a 0:node where mass or charge is conserved. The reason to lump these two together is that mass or charge conservation always includes a local storage term, so the storage term can be thought of as internal to the 0:node. Combining them in this way of course, greatly simplifies the diagram for the bond graph model of a complex system.

Another convention adopted here is to use a *superscript* to indicate the electrical or mechanical quantity or chemical species being expressed, and a *subscript* to indicate the location of that quantity (e.g. in the extracellular or intracellular fluid, or the cell membrane, etc).

The flux balance equations associated with the 0-nodes are:

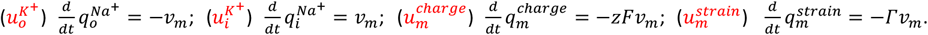

*z* = 1 for *Na*^+^, and *F* has units C.mol^-1^ (to convert the molar flux *v*_*m*_ to a charge flux). *Γ* has units mol^-1^ (to convert the molar flux *v*_*m*_ to a strain rate).

The energy balance equations associated with the 1-nodes are:

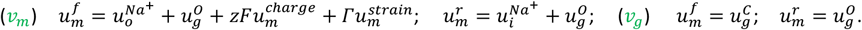

The reaction fluxes *v*_*m*_ (mol.s^-1^) and *v*_*a*_ (mol.s^-1^) are

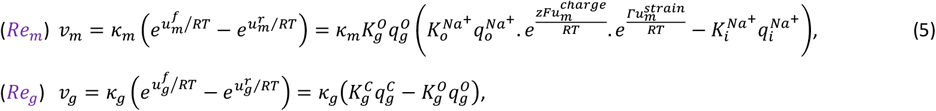

where the second step in both cases uses the expressions for the forward and reverse potentials inserted into the energy balance equations.

Note that *Γ* has units of mol^-1^, consistent with a power balance 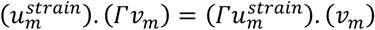 where (since strain is dimensionless) power (J.s^-1^) on the left has units (J).(mol^-1^.mol.s^-1^) and power on the right has units (mol^-1^.J).(mol.s^-1^).

Since the total number of gates 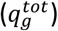 is constant 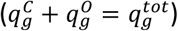, the probabilities of the gates being open and closed, respectively, are

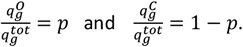

where (0 ≤ *p* ≤ 1).

The gate mass balance equations 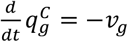 and 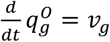 are therefore represented by

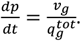

Using the bond graph flux equation 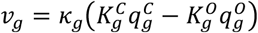, gives

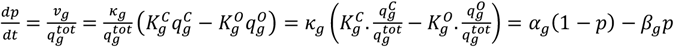

where 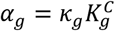 is the rate at which closed gates open, and 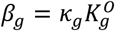 is the rate at which open gates close. For voltage gated ion channels, these opening and closing rate constants are defined as functions of the membrane potential 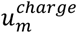. Equation 5 now becomes

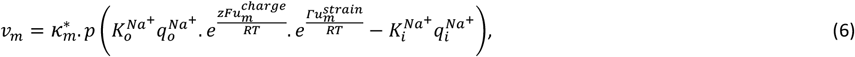

where 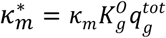 is the open channel conductance.

With the molar flux given by *v*_*m*_, the electrical current flow through the membrane is

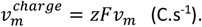

The chemical flux *v* (mol.s^-1^) or electrical current flow 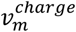 (C.s^-1^) is zero at the equilibrium or ‘Nernst’ potential,

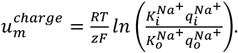

The concentration of species are given by 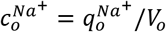 and 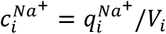. Therefore, since the thermodynamic constants are related by 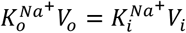,

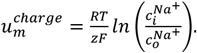

This is the form of the Nernst equation usually used in electrophysiology. But see [8] for an alternative constant-field Goldman-Hodgkin-Katz (GHK) model of ion permeation that accounts for ion channel rectification.

#### Example 3: An enzyme-catalyzed reaction

Now consider the enzymatic reaction shown in Figure 5a, which is often associated with Michaelis-Menten (MM) kinetics [9]. 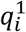 is a substrate that binds reversibly to an enzyme 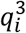 to form the complex 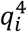, which breaks down to regenerate the enzyme and yield a product 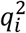. In conventional MM kinetics this last step is treated as irreversible, if 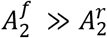.

**Figure 5.**
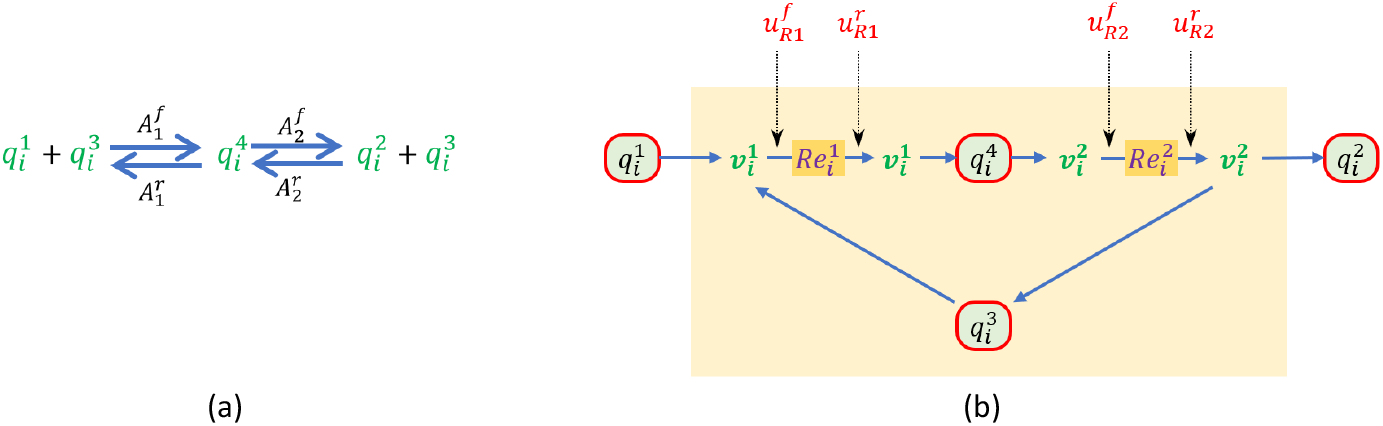
(a) An enzyme 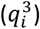-catalysed reaction, and (b) its bond graph representation. Flux balance is ensured for each of the four species at the 0:nodes, and energy balance at the 1:nodes ensures the correct stoichiometry. The forward and reverse potentials, for each of the two reactions, are indicated by the dotted arrows.

The flux balance equations for the four species, defined at the four 0:nodes, are

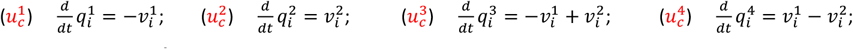

Note that since 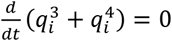, the total amount of enzyme (including in its bound form 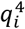) is constant. i.e.

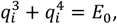

where *E*_0_ is the initial quantity of enzyme.

The energy balance equations, defined at the two 1:nodes, are

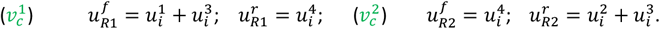

Using these potentials, the two reactions are

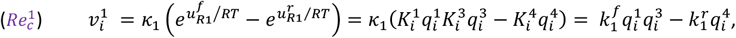

where 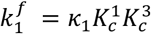 (mol^-1^.s^-1^) and 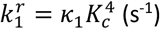, and

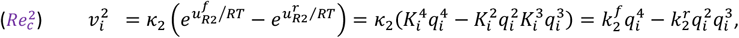

where 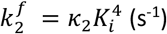 and 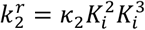 (mol^-1^.s^-1^).

The *Briggs-Haldane* analysis of the reaction [7] assumes that there is a much higher concentration of substrate than enzyme 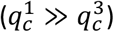 and that the complex 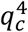 therefore quickly reaches a steady-state (SS).

Assuming a steady constant flux with 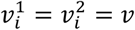 and 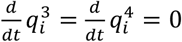,

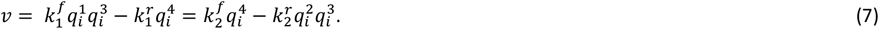

From the second equation in (7) we can express 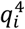 in terms of 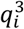:

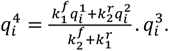

Using conservation of total enzyme 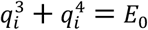,

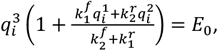

which, with equation 7, gives

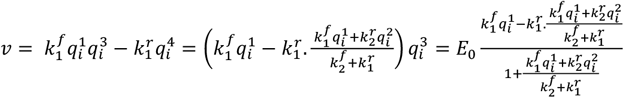

Or

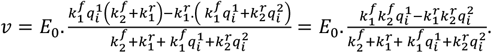

Substituting back 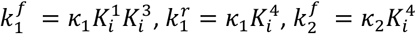 and 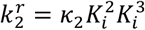, gives

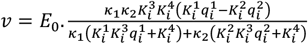

or, rearranging the denominator,

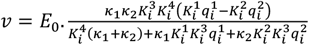

A useful way of expressing this relationship between the SS flux and the solute quantities is

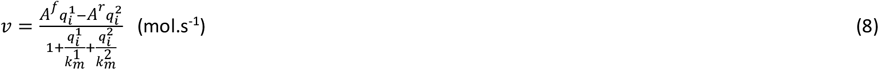

where

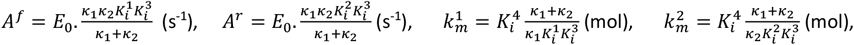

since this highlights the relationship with the Michaelis-Menten (MM) flux expression below.

Note that since 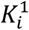 (mol^-1^) and 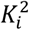 (mol^-1^) are the thermodynamic constants associated with the solute (not the reaction), the reaction flux is defined by three combinations of biophysical parameters 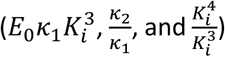 – one more than the MM flux expression below, and since 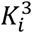 and 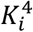 are usually assumed to be the same, this reduces to only two parameters needed for fitting experimental data.

The MM approximation goes one step further and assumes that with a sufficiently low concentration of product 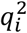 relative to the complex, the 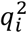 term in equation 8 can be ignored, and

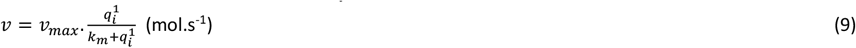

where

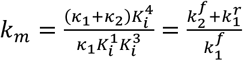 is the MM constant and 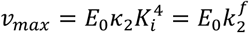 is the maximal (SS) flux.

Note, however, that the MM approximation assumes that the reaction is irreversible, which violates thermodynamic principles.

With the above background to bond graph modelling of physical (including physiological) processes, we can now use this approach to derive the equations governing SLC transporters, using *SLC2A2* and *SLC5A1* as specific examples from which more general lessons can be derived for the entire family.

## RESULTS

### The SLC superfamily

The SLC superfamily currently includes 62 families of SLC transporters [1] that deal with the transport of the following small molecules:

Cations: 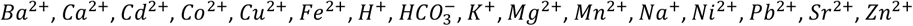, ammonium 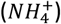 Anions: *Cl*^−^, bicarbonate 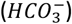, phosphate 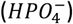, pyruvate 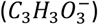,

Amino acids: Ala, Arg, Asn, Asp, Cys, Gln, Glu, Gly, His, Ise, Leu, Lys, Met, Phe, Pro, Ser, Thr, Trp, Tyr, Val

Sugars: glucose (Glc),

Hormones & neurotransmitters: acetylcholine (ACh), epinephrine, norepinephrine (NE), steroids

Vitamins: folate (B9), pyridoxine (B6), thiamine (B1),

Lipids: cholesterol, sphingosine,

Others: bile acids, heme, selenate, sulfate, thiosulfate, riboflavin, molybdate, pyrophosphate (*H*_4_*P*_2_*O*_7_)

We briefly describe some features of these transporters before looking in detail at members of two families that are involved in transporting glucose across cell membranes: *SLC2A2* and *SLC5A1*.

Table 1 lists the members of the first family (*SLC1*), together with their familiar protein name, their UniProt IDs, the substrate(s) carried by the transporter and a diagram of the chemistry.

**Table 1.**
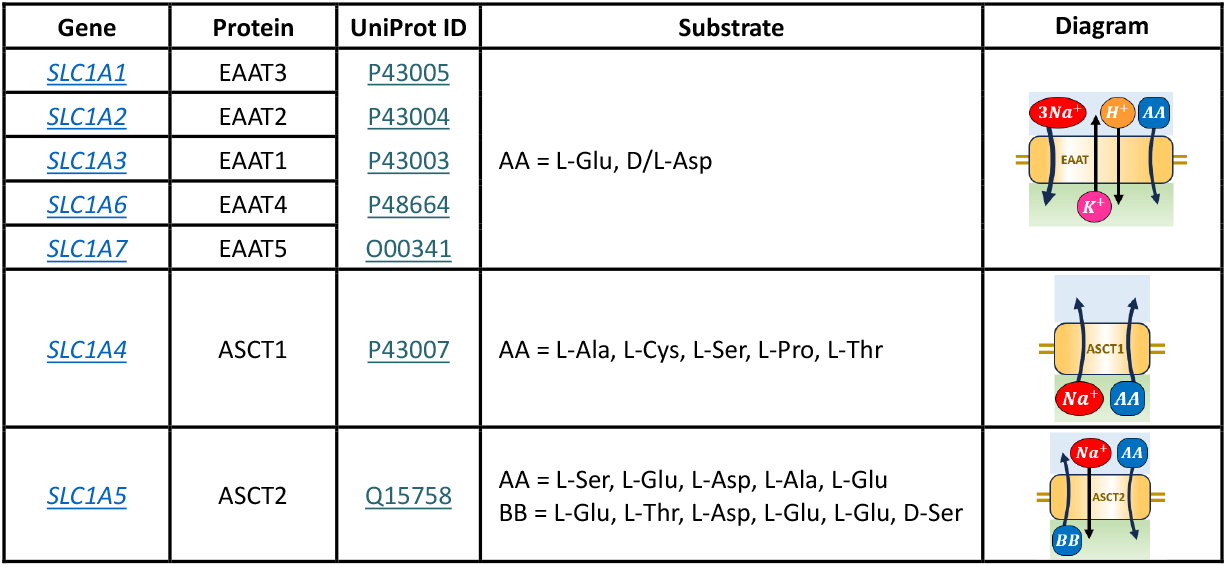
The first family (SLC1) in the SLC superfamily. Note that the extracellular space is shown above the bilipid membrane and the intracellular space below the membrane in the diagrams on the right.

### Facilitated diffusion with *SLC2A2* (GLUT2)

The review article [10] provides a comprehensive overview of SLC2 family of transporters. While alternative models for SLC2A2 (GLUT2) were proposed to address inconsistent observations in some experiments [11], most kinetic and biophysical data support the alternating conformation mechanism of SLC2A2 (GLUT2) transporter [12, 13, 14]. There has been debate over whether the alternating models violate the energy conservation laws [15, 16]. This paper uses the most accepted alternating model [12, 13, 14] to demonstrate that the bond graph approach describes the energetic perspectives of a system in a more explicit manner.

In this section we present a modelling pipeline for the *SLC2A2* (GLUT2) facilitated diffusion of glucose through a bilipid membrane (see Figure 6). The pipeline goes from (a) the statement of the biochemical reaction, to (b) a bond graph diagram of the full kinetics of the transport process, to (c) the steady-state flux model. We demonstrate parameter fitting for both the full kinetic model using the biophysical parameters and the reduced steady-state flux model using both the full set of biophysical parameters and a reduced set of empirical parameters.

**Figure 6.**
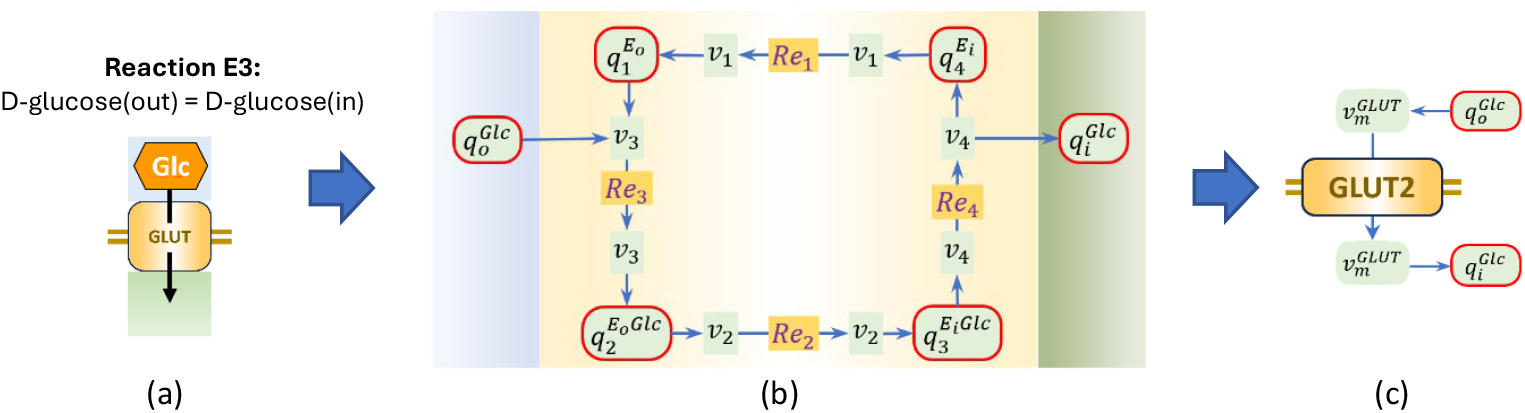
The pipeline from (a) the chemical reaction with its representative icon, (b) the bond graph diagram for the reaction, and (c) the diagram for the reduced model showing the steady-state flux dependencies on the molar quantities 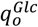 and 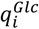.

The reactions represented by Figure 6b are as follows:

1. *Re*_1_: the transition of the protein from the inward-facing state to an outward-facing state;
2. *Re*_3_: the binding of the ligand (external glucose 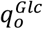) to the outward-facing protein;
3. *Re*_2_: the transition of the protein from the outward-facing state to an inward-facing state;
4. *Re*_4_: the unbinding of the ligand (external glucose 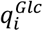) from the inward-facing protein;

In the following equations glucose (*Glc*) is represented by the symbol *A* as a generic (uncharged) ligand since these equations are valid for facilitated diffusion of any electrically neutral molecule across a membrane.

The flux balance equations associated with the 0-nodes are:

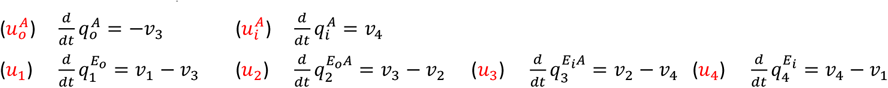

The energy balance equations associated with the 1-nodes are:

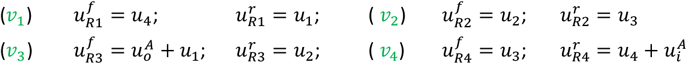

The constitutive laws for the storage terms are:

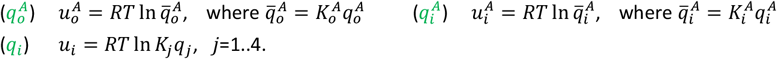

Note that we nondimensionalise the solute quantities (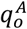 and 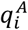) by using 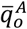 and 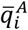, but retain the thermodynamic constants (*K*_1_, *K*_2_, *K*_3_, *K*_4_,) for the protein state variables as these quantities (*q*_*j*_) must sum to a constant total (*q*_*tot*_) – see below.

The reactions, with the substituted potentials, are:

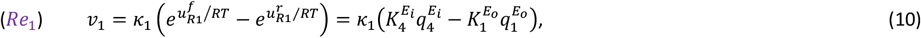

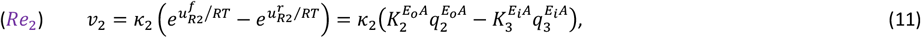

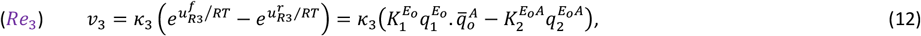

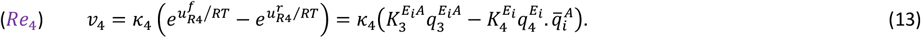

Conservation of the enzyme requires the constraint that

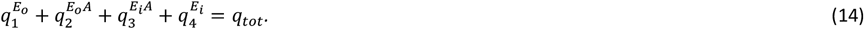

The flux balance equations, with the flux terms from equations 10..13 and the enzyme conservation equation 14, can be solved for the 6 molar quantities (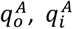 and 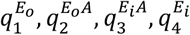) subject to appropriate initial conditions, as illustrated below for specific experimental conditions.

However, before we use experimental data to fit the 8 protein parameters (4 thermodynamic constants and 4 reaction rates) of this full kinetic bond graph model, we consider the steady-state situation, which yields an analytic expression for the flux as a function of the solute quantities.

#### Facilitated diffusion with steady-state flux and rapid binding and unbinding

A high ratio of substrate to enzyme (the Briggs-Haldane assumption) implies steady-state conditions:

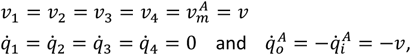

(dropping the superscripts on the protein states).

We also assume that the binding and unbinding rates for the solute molecule are much faster than the carrier state transition rates [13], in which case (*k*_3_, *k*_4_ → ∞), the bracketed terms on the RHS of (4.1.3) and (4.1.4) must be zero, and therefore

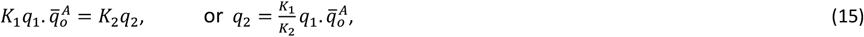

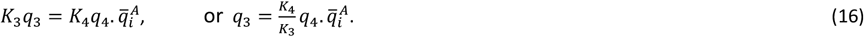

Substituting *q*_2_ and *q*_3_ into the other two reactions, assuming steady-state with *v*_1_ = *v*_2_ = *v*, gives

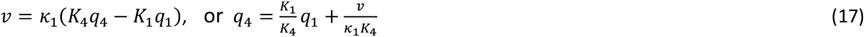

and

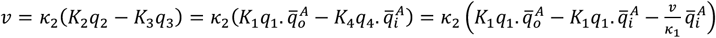

from which

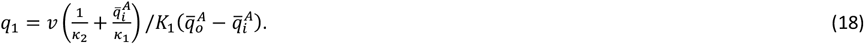

Substituting (15), (16), (17) into (14), gives

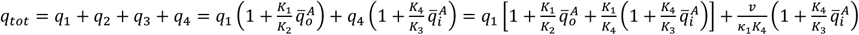

with *q*_1_ given by (18). i.e.,

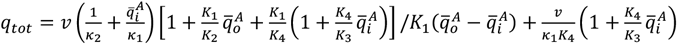

Or

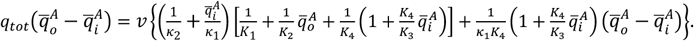

Rearranging for *v*,

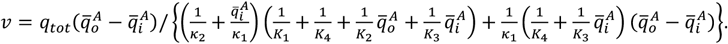

Multiplying numerator and denominator by *k*_1_*K*_3_

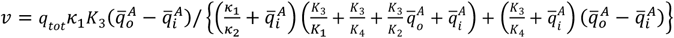

or, rearranging the denominator,

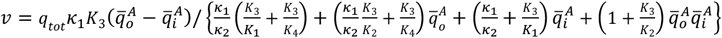

or

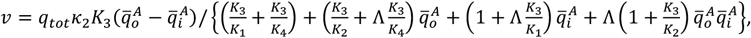

where 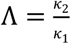 is the forward to reverse ratio of enzyme state transitions.

It is convenient to express this relationship as

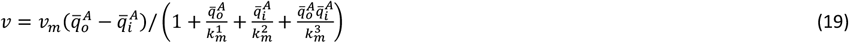

Where

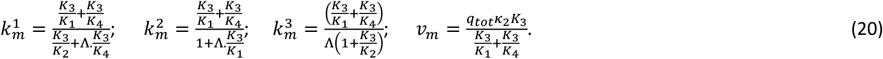

Note that 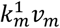 is the maximum flux obtained as 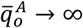 with 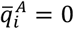, and 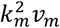 is the maximum flux obtained as 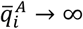 with 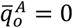, as shown in Figure 7. All the 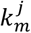 (j=1.3) terms are dimensionless.

**Figure 7.**
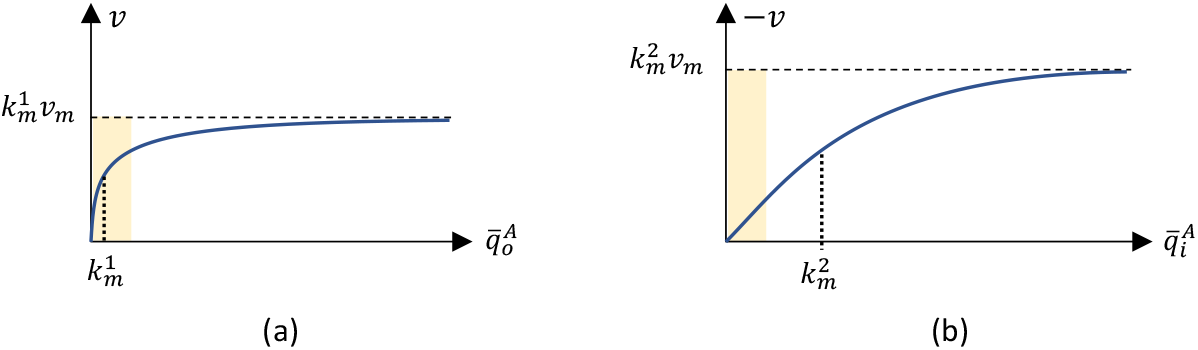
(a) Inward flux as a function of 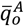 when 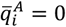; (b) Outward flux as a function of 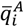. when 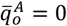. Typical operating ranges for 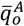 and 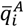 are shown by the shaded blocks. Notice that a relatively low value for 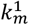 (compared with 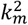) ensures that the inward flux is relatively much higher than the outward flux. Removal of 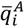 from the intracellular environment, due to its involvement in other reactions, also keeps 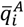 low.

#### Fitting the bond graph model of facilitated diffusion to experimental observations

The full kinetic model for facilitated diffusion is given by the 6 flux balance equations, the 4 flux expressions (10 to 13) and the mass constraint equation 14. Here we use experimental kinetic data from the literature [13] to fit the 9 biophysical parameters (the 4 reaction rate constants *k*_1_ to *k*_4_, the 4 thermodynamic constants *K*_1_ to *K*_4_ and the total amount of enzyme *q*_*tot*_) in those equations.

The kinetic parameters from [13] for each reaction in Figure 6(b) are listed in Table 2(a). The kinetic parameters of reactions *Re*_3_ and *Re*_4_, which are not given in [13], are set to arbitrarily large numbers to align with the fast binding assumptions while applying the constraints (defined in [13]): 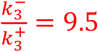 (mM) and 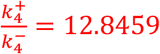 (mM). We applied the method introduced in [17] to convert the thermodynamically consistent kinetic parameters in Table 2(a) to the bond graph parameters in Table 2(b). The detailed fitting process and parameters can be found in the accompanying *Physiome* paper [18]. We simulate the full bond graph model using the fitted parameters and the steady-state predictions for the full kinetic model are shown in Figure 8 as black lines.

**Table 2.** (a) The biophysical parameters for the full bond graph model of SLC2A2, and (b) the empirical parameters of the reduced steady-state model given by Equation 19 derived from the bond graph model.

| Reaction | Forward rate constant $k_i^+$ | Reverse rate constant $k_i^-$ | Ref |
| --- | --- | --- | --- |
| $Re_1$ | $0.726 \text{ s}^{-1}$ | $12.1 \text{ s}^{-1}$ | [13] |
| $Re_2$ | $1113 \text{ s}^{-1}$ | $90.3 \text{ s}^{-1}$ | [13] |
| $Re_3$ | $4.5e7 \text{ mM}^{-1} \text{ s}^{-1}$ | $4.5e7 \times 9.5 \text{ s}^{-1}$ | Not given in [13] |
| $Re_4$ | $2.7e5 \times 12.8459 \text{ s}^{-1}$ | $2.7e5 \text{ mM}^{-1} \text{ s}^{-1}$ | Not given in [13] |

(a)
| Parameters | Value | Unit |
| --- | --- | --- |
| $K_i^A$ | 149.65 | $\text{fmol}^{-1}$ |
| $K_o^A$ | 149.65 | $\text{fmol}^{-1}$ |
| $K_1$ | 33.20 | $\text{fmol}^{-1}$ |
| $K_2$ | 4.25e3 | $\text{fmol}^{-1}$ |
| $K_3$ | 344.59 | $\text{fmol}^{-1}$ |
| $K_4$ | 1.99 | $\text{fmol}^{-1}$ |
| $\kappa_1$ | 0.36 | $\text{fmol.s}^{-1}$ |
| $\kappa_2$ | 0.26 | $\text{fmol.s}^{-1}$ |
| $\kappa_3$ | 1.01e5 | $\text{fmol.s}^{-1}$ |
| $\kappa_4$ | 1.01e4 | $\text{fmol.s}^{-1}$ |

(b)
| Parameters | Value | Unit |
| --- | --- | --- |
| $v_m$ | 0.003284 | $\text{fmol.s}^{-1}$ |
| $k_m^1$ | 1.4735 | dimensionless |
| $k_m^2$ | 21.671 | dimensionless |
| $k_m^3$ | 235.07 | dimensionless |

**Figure 8.**
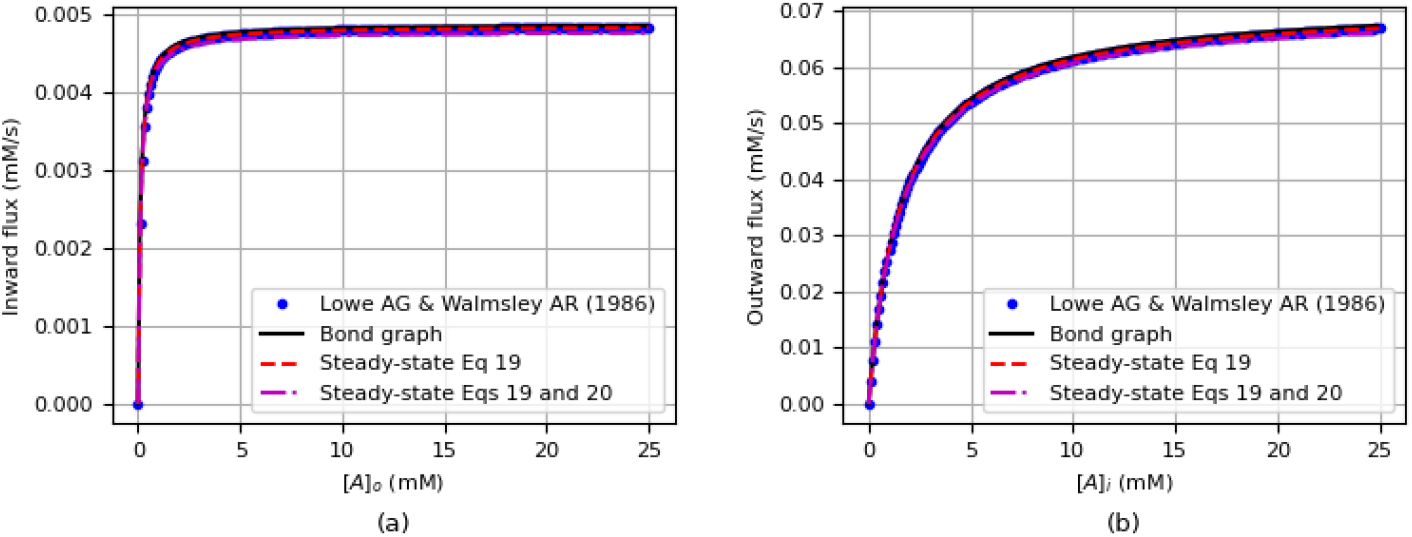
(a) Inward flux as a function of [*A*]_*o*_ when [*A*]_*i*_ = 0, and (b) outward flux as a function of [*A*]_*i*_ when [*A*]_*o*_ = 0. Note that in order to compare with the kinetic data in Low AG & Walmsley AR (1986) [13], the molar amount of glucose in the bond graph model was converted to glucose concentrations. The inward flux in Low AG & Walmsley AR (1986) was computed using 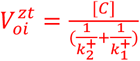, while the outward flux was calculated by 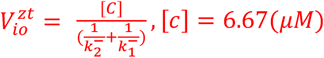. The information needed to reproduce these results can be found in [18].

Equation 19 gives the SS flux under the assumption that binding and unbinding occur very rapidly in comparison with the transition rates for the carrier protein and that the enzyme is cycling at a constant rate. We also show here how the 4 parameters 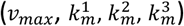 in Equation 19 can be fitted directly with steady-state flux data and that these values match the predictions (Equation 20) determined by the parameters fitted to the full kinetic model.

### Steady-state flux experiments

In their first experiment, Lowe and Walmsley [13] set the intracellular concentration to 0 (mM) and measured the inward flux of glucose for a varying range of extracellular glucose concentration.

Note that the concentration of glucose carrier molecules in human red blood cells was observed to be 6.67×10^−3^ mM [13].

Putting 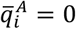 in Equation 19 with

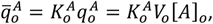

where [*A*]_*o*_ (mM or mol.m^-3^) is the concentration of glucose, and *V*_*o*_ (m^3^) is the volume of the extracellular compartment, gives the inward flux *v*_*oi*_ (‘oi’=outside→inside) as

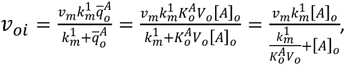

Or

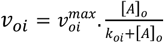

where 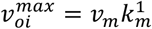 (mM.s^-1^) is the maximum flux (as [*A*] → ∞), and 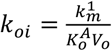 (mM) is the concentration at which 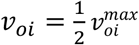. Note that specifying the parameter 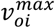 in units of mM.s^-1^, means that *v*_*oi*_ also has units of mM.s^-1^. The maximum flux observed experimentally in [13] is 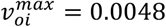 (mM.s^-1^) and the fitted Michaelis constant is *k*_*oi*_= 0.1094 (mM), so

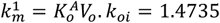

using *V*_*o*_ = 0.09 (*pL*) and 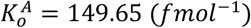, (see [18], for details of fitting procedures).

In their second experiment, Lowe and Walmsley [13] set the extracellular concentration to 0 (mM) and measured the outward flux of glucose *v*_*io*_ at varying intracellular glucose concentration [*A*]_*i*_.

Again, by comparing Equation 19 with the Michaelis-Menten graph of zero trans influx given by [13], 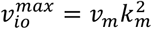 (mM.s^-1^) (the maximum flux (as [*A*]_*i*_ → ∞), and 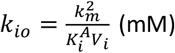 (the concentration at which 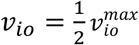. This gives 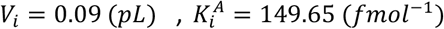 and *k*_*io*_ = 1.609 (*mM*), and hence

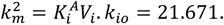

The red dashed lines in Figure 8 show the predictions of the steady-state model (Equation 19) using these directly fitted parameters.

The parameter values found for the full bond graph model and the reduced steady-state model are given in Table 2(b) and (c), respectively. Notice that *k*^3^, which weights the product term in equation 19, is an order of magnitude higher than the other two *k*_*m*_ parameters and indicates that this term contributes very little to the flux.

The magenta lines in Figure 8 show the steady-state model predictions using the empirical parameters defined in Equation 20 using the physical parameters that are listed in Table 2(b):

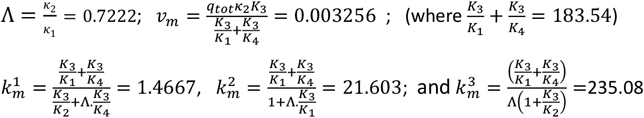

Note how closely these match the fitted empirical parameters listed in Table 2(c).

From Figure 8, we can see that both the steady-state prediction by the full bond graph model and the steady-state Equation 19 derived from the bond graph model can accurately reproduce the influxes and effluxes from the data of [13].

### Sodium-glucose cotransport with *SLC5A1* (SGLT1)

The six-state energetic kinetic model of sodium-glucose cotransporter SLC5A1 (SGLT1) was proposed by Parent et al. [19] to account for most experimental observations [20, 21, 22]. A simplified electroneutral bond graph description of the SGLT1 model has been presented in [23], while the electrogenic version is available in [24] with different parameters and bond graph formulations. Here, we use this widely adopted example with the original kinetic parameters in [19] to show how the proposed modelling pipeline can handle multiple physical domains (electrical and chemical) in a unified framework.

The SGLT1 (*SLC5A1*) sodium-driven transport of glucose is shown in Figure 9. The equations from the bond graph diagram and the calculation of the steady-state flux are given below.

**Figure 9.**
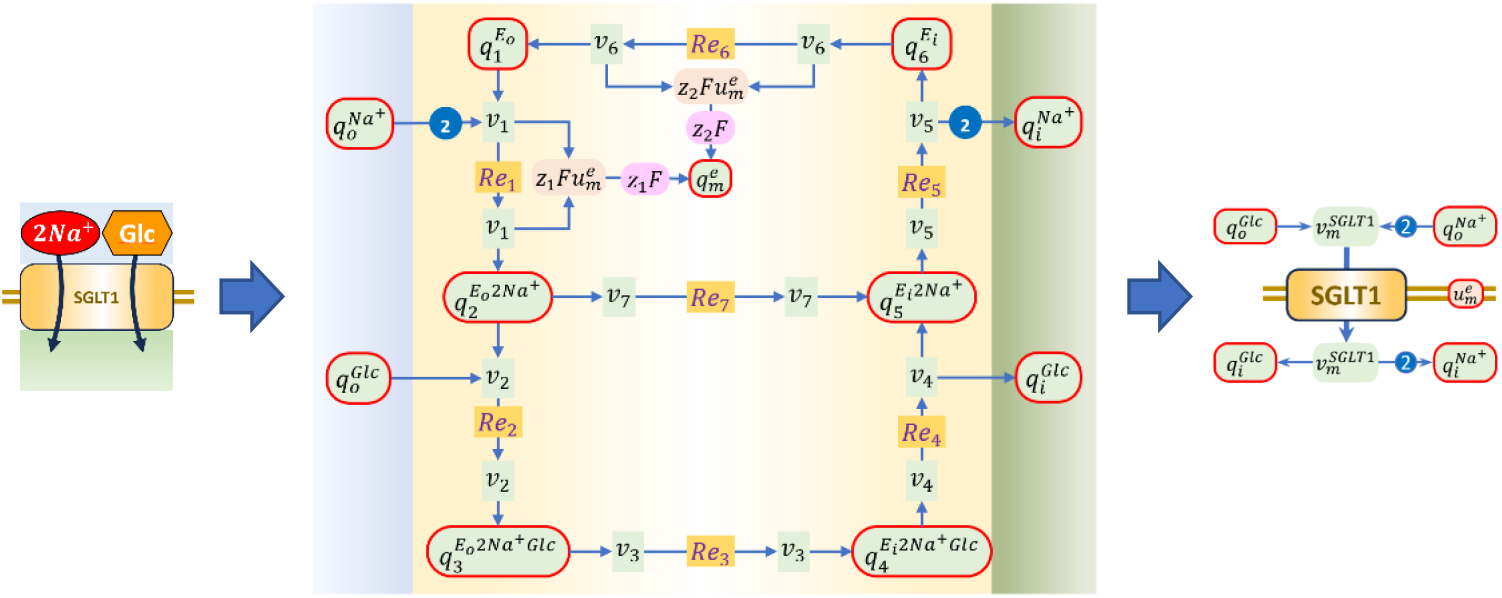
The modelling pipeline for SLC5A1. Note the addition of the links with the transmembrane potential in this electrogenic reaction, and the use of the blue symbol showing the number of moles of *Na*^+^ entering or exiting the reaction, per mole of reaction flux 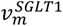 crossing the membrane. The reduced (steady-state) form of the model is shown on the right.

The flux balance equations associated with the 0-nodes are:

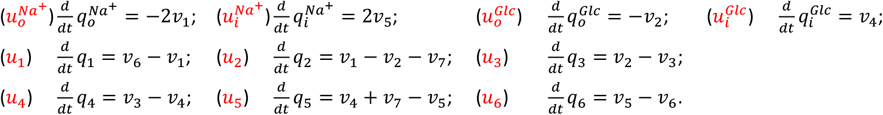

These differential equations provide the flux balance at the ten 0:nodes indicated in Figure 9 by the red-encircled storage quantities.

The energy balance equations associated with the 1-nodes are:

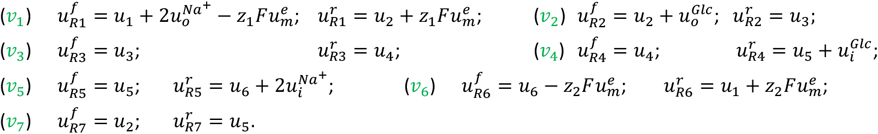

These algebraic stoichiometry equations provide energy balance for the seven fluxes shown by the *v*_*i*_ terms in Figure 9. Note that charge transfer is also modeled.

The seven reactions in Figure 9 give rise to the following fluxes:

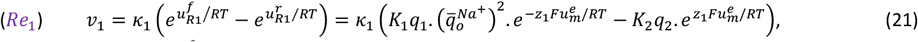

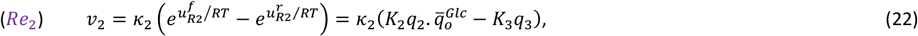

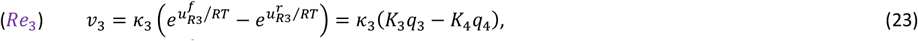

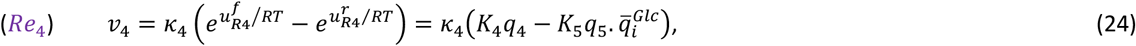

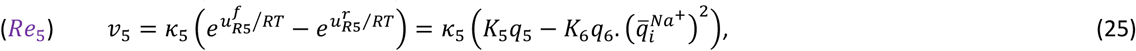

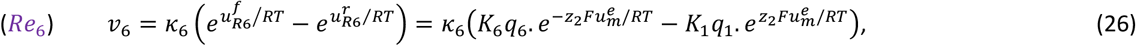

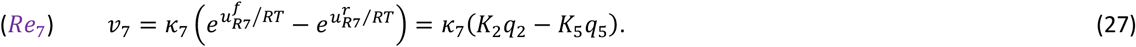

These equations are supplemented with the constraint on the total amount of protein (*q*_*tot*_):

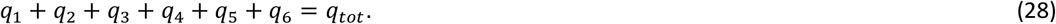

By summing up potentials, the overall affinity of the transporter cycle is

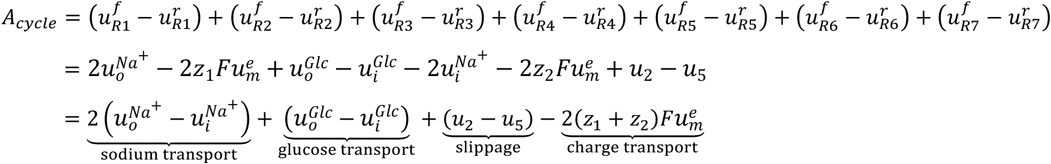

Reaction 7 represents the possibility for the transporter to transition from outward facing to inward facing without binding glucose (called ‘slippage’). Since the transporter cycle moves 2 units of charge into the cell per cycle, *z*_1_ + *z*_2_ = 1, and (ignoring slippage)

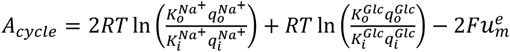

At equilibrium, *A*_*cycle*_ = 0 and the transporter stalls. The reversal potential is the membrane potential

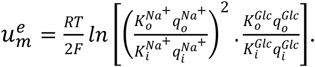

Solving the 10 flux balance equations using the fluxes from the 7 reactions (Equation 21 to Equation 27), together with the enzyme mass constraint (Equation 28) and specified values for 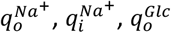 and 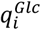 gives 7 equations in the 7 variables *q*_1_ to *q*_6_ and *q*_*tot*_. These equations include 14 biophysical parameters (the 6 thermodynamic parameters *K*_1_ to *K*_6_, the 7 reaction rate parameters *k*_1_ to *k*_7_, and the total amount of enzyme *q*_*tot*_). These parameters are fitted to experimental data from Parent et al. [19] below.

#### Sodium-glucose cotransport with steady-state flux and rapid binding and unbinding

To derive an analytic formula for the steady-state behaviour of the transporter, we make the usual two assumptions: (i) that the substrates are present in much higher quantities than the membrane-bound transporter (the Briggs-Haldane assumption), and that the cycle is therefore transitioning through the 6 states at a constant steady-state rate *v*, and (ii) that the binding and unbinding reactions (1, 2, 4 and 5) are much faster than the state transitions between inward- and outward-facing states of the protein. We make the further assumption that slippage can be ignored (*v*_7_ = 0).

From the first of these,

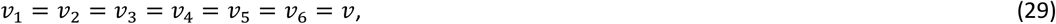

and from the second,

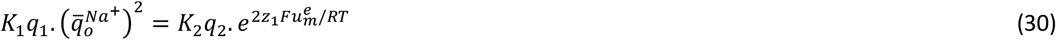

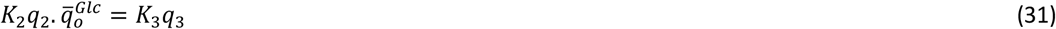

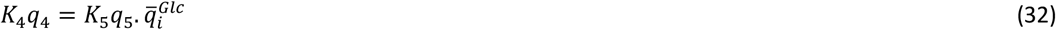

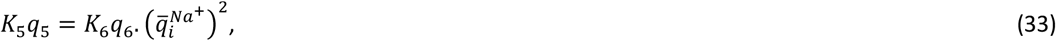

leaving *v*_3_ = *v*_6_ = *v*, or

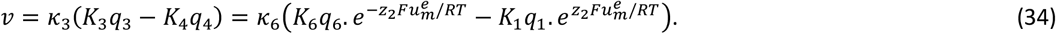

The 6 protein states *q*_1_.. *q*_6_ can be eliminated from equations 30 to 34 and 28, to yield an expression for the flux *v* in terms of the 4 solutes 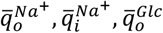 and 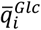. From 30 and 31,

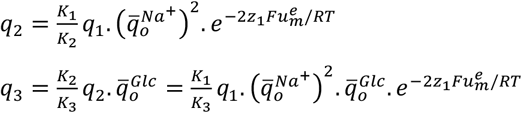

From 32 and 33,

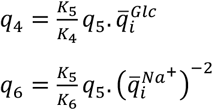

Substituting the last 3 equations into the second equation of 34,

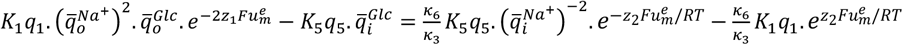

which gives

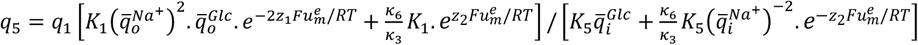

Or

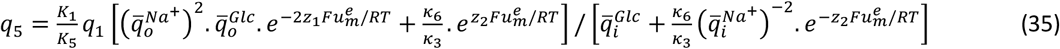

Substituting for *q*_2_.. *q*_6_ in terms of *q*_1_ in Equation 28, gives

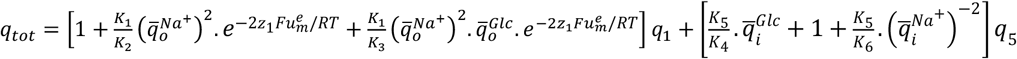

or, using *q*_5_ given in terms of *q*_1_ by Equation 35:

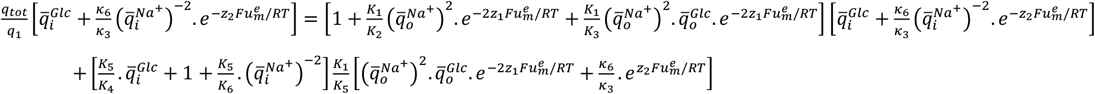

Or

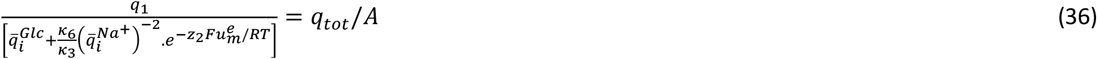

Where

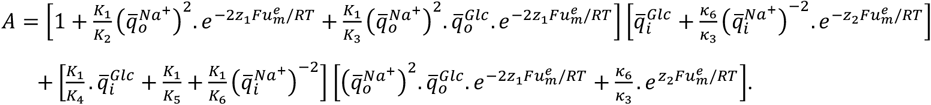

From the first equation in (34),

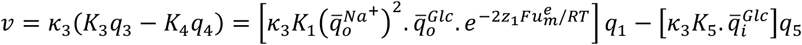

Or

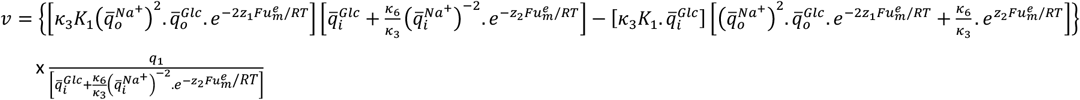

Or

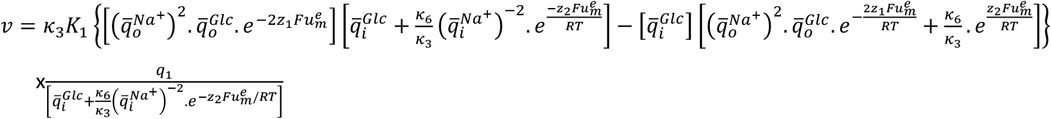

Or

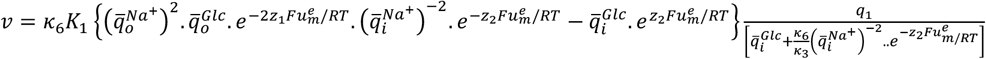

or, now substituting for *q*_1_ from Equation 36,

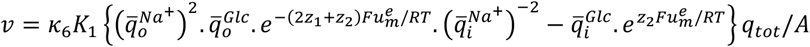

or, multiplying numerator and denominator by 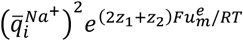

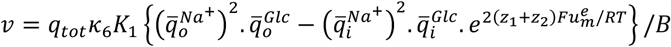

Where

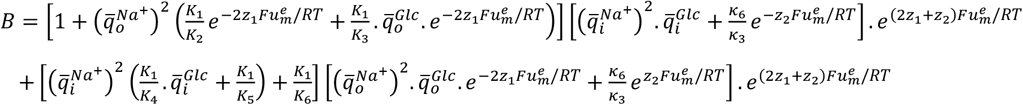

where *z*_1_ + *z*_2_ = 1.

Hence

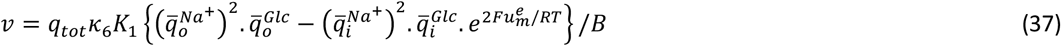

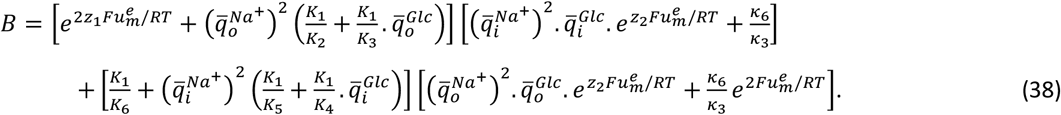

Note that the flux is zero when

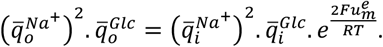

i.e. the equilibrium potential is, as above,

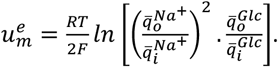

In Figure 10 the current-voltage (I-V) relationship for the full kinetic bond graph model (run to steady state), and the reduced steady-state flux model given by equations 37 and 38, are compared using biophysical parameters fitted to experimental data from Parent et al. [19] (see below) but with *k*_1_, *k*_2_, *k*_4_ and *k*_5_ set to arbitrarily high values to reflect the fast binding and unbinding assumption. This result confirms that the assumption of no slippage is valid for the range of potentials shown (the slight discrepancy at the lower voltages is due to this small slippage flux).

**Figure 10.**
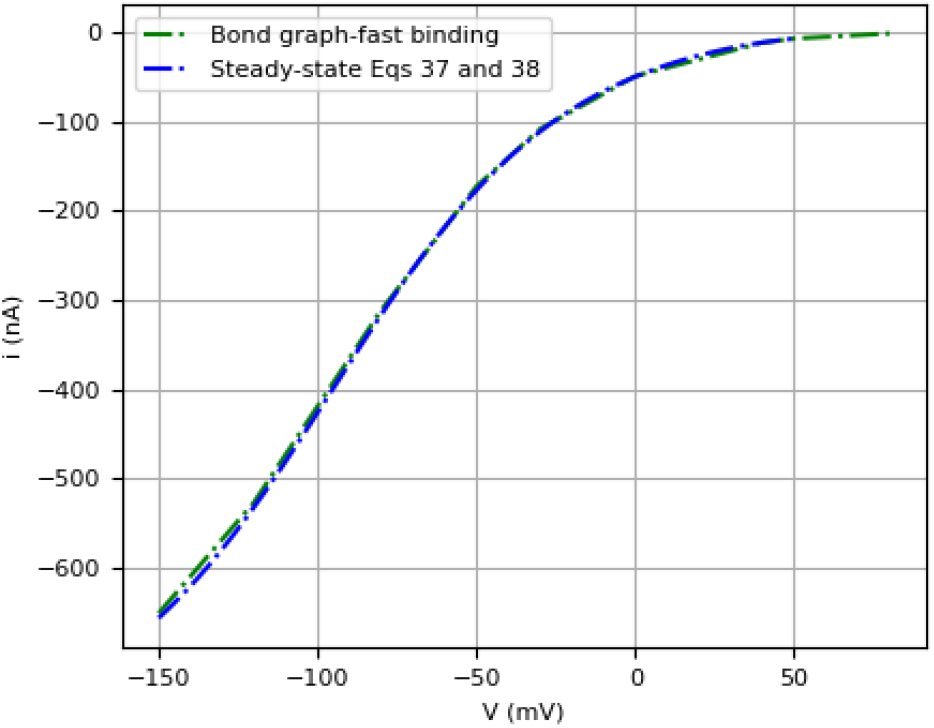
The steady-state results predicted by the full bond graph model, compared with the results from the reduced steady-state model. Both simulations use the assumption of fast binding and unbinding. The information needed to reproduce these results can be found in [18].

#### Fitting the bond graph model of sodium-glucose cotransport to experimental observations

The kinetic parameters [19] of each reaction in Figure 9(b) are listed in Table 3(a). The reverse rate constant 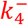 of reaction *Re*_4_ and 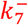 of reaction *Re*_7_ are calculated by the detailed balance equations:

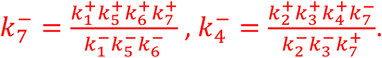

**Table 3.** (a) The thermodynamic parameters and (b) reaction rate constants for the bond graph model of *SLC5A1* shown in Figure 9. Note the very small rate constant (*k*_7_) for slippage. The kinetic parameter 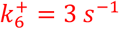 was used in the steady-state plot in [19]. This value and the corresponding bond graph parameters are in brackets ().

| Reaction | Forward rate constant $k_i^+$ | Reverse rate constant $k_i^-$ | Ref |
| --- | --- | --- | --- |
| $Re_1$ | $80000 M^{-2} s^{-1}$ | $500 s^{-1}$ | [19] |
| $Re_2$ | $1e5 M^{-1} s^{-1}$ | $20 s^{-1}$ | [19] |
| $Re_3$ | $50 s^{-1}$ | $50 s^{-1}$ | [19] |
| $Re_4$ | $800 s^{-1}$ | $1.8285e7 M^{-1} s^{-1}$ ($1.0971e7 M^{-1} s^{-1}$) | $k_4^-$ is calculated by the detailed balance |
| $Re_5$ | $10 s^{-1}$ | $50 M^{-2} s^{-1}$ | [19] |
| $Re_6$ | $5 s^{-1}$ ($3 s^{-1}$) | $35 s^{-1}$ | [19] |
| $Re_7$ | $0.3 s^{-1}$ | $1.371 s^{-1}$ ($0.823 s^{-1}$) | $k_7^-$ is calculated by the detailed balance |

(a)
| Parameters | Value | Unit |
| --- | --- | --- |
| $K_i^{Na}$ | 3.22e-8 | fmol<sup>-1</sup> |
| $K_o^{Na}$ | 3.22e-8 | fmol<sup>-1</sup> |
| $K_i^{Glc}$ | 4.85e-6 | fmol<sup>-1</sup> |
| $K_o^{Glc}$ | 4.85e-6 | fmol<sup>-1</sup> |
| $K_1$ | 2.235 | fmol<sup>-1</sup> |
| $K_2$ | 10.44 | fmol<sup>-1</sup> |
| $K_3$ | 8.602 | fmol<sup>-1</sup> |
| $K_4$ | 8.602 | fmol<sup>-1</sup> |
| $K_5$ | 47.71 (28.63) | fmol<sup>-1</sup> |
| $K_6$ | 0.319 (0.192) | fmol<sup>-1</sup> |

(b)
| Parameters | Value | Unit |
| --- | --- | --- |
| $\kappa_1$ | 47.91 | fmol.s<sup>-1</sup> |
| $\kappa_2$ | 2.325 | fmol.s<sup>-1</sup> |
| $\kappa_3$ | 5.812 | fmol.s<sup>-1</sup> |
| $\kappa_4$ | 93 | fmol.s<sup>-1</sup> |
| $\kappa_5$ | 0.21 (0.349) | fmol.s<sup>-1</sup> |
| $\kappa_6$ | 15.66 | fmol.s<sup>-1</sup> |
| $\kappa_7$ | 0.029 | fmol.s<sup>-1</sup> |

Similarly, we applied the method introduced in [17] to convert the thermodynamically consistent kinetic parameters in Table 3(a) to the bond graph parameters in Table 3(b) and Table 3(c).

Figure 11 shows the results of fitting the 14 biophysical parameters of the full bond graph model to transient electrical current measurements by Parent et al. [19] at clamped membrane voltages of 50mV and -150mV. Figure 11(a) shows the experimental results and model predictions for the case when the external glucose level is set to zero, and Figure 11(b) shows the measured results and model predictions for the case when the external glucose level is set to 1mM.

**Figure 11.**
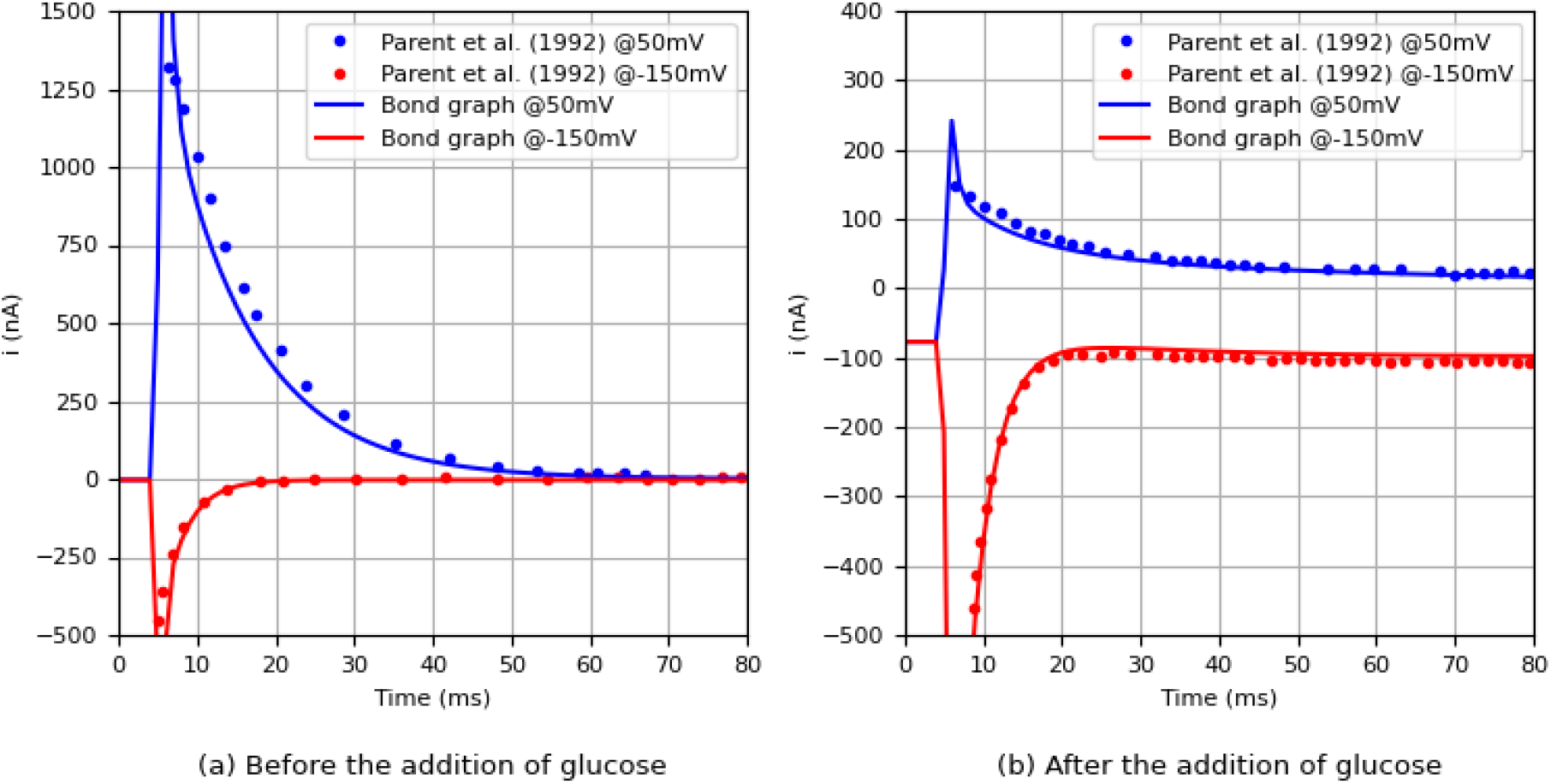
The time course of the carrier-mediated currents. (a) The electrical current when [*Glc*]_*o*_ = 0 mM, and (b) the current when [*Glc*]_*o*_ = 1 mM; The output of the bond graph model is the current −*I*_*i*_, and the data of Parent et al. [19] are reproduced from Figure 10 of that paper using the digitizing software Engauge. The information needed to reproduce these results can be found in [18].

We applied a range of test potentials to the full bond graph model with a slightly reduced 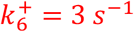 to produce the steady-state glucose-dependent I-V curve (red plot) shown in Figure 12, which is compared with the I-V curve (black dot plot) given in Figure 5 in [19]. The simulated glucose-dependent current is defined as the difference in the carrier-mediated current at steady state before and after the addition of glucose. As noted in [19], the background current induced by the experimental conditions can be accounted for by an additional RC circuit. However, this element is not included in the bond graph model, which may explain the discrepancy between the simulation and the measurements. Additionally, errors may be introduced during the digitizing process of the published figure.

**Figure 12.**
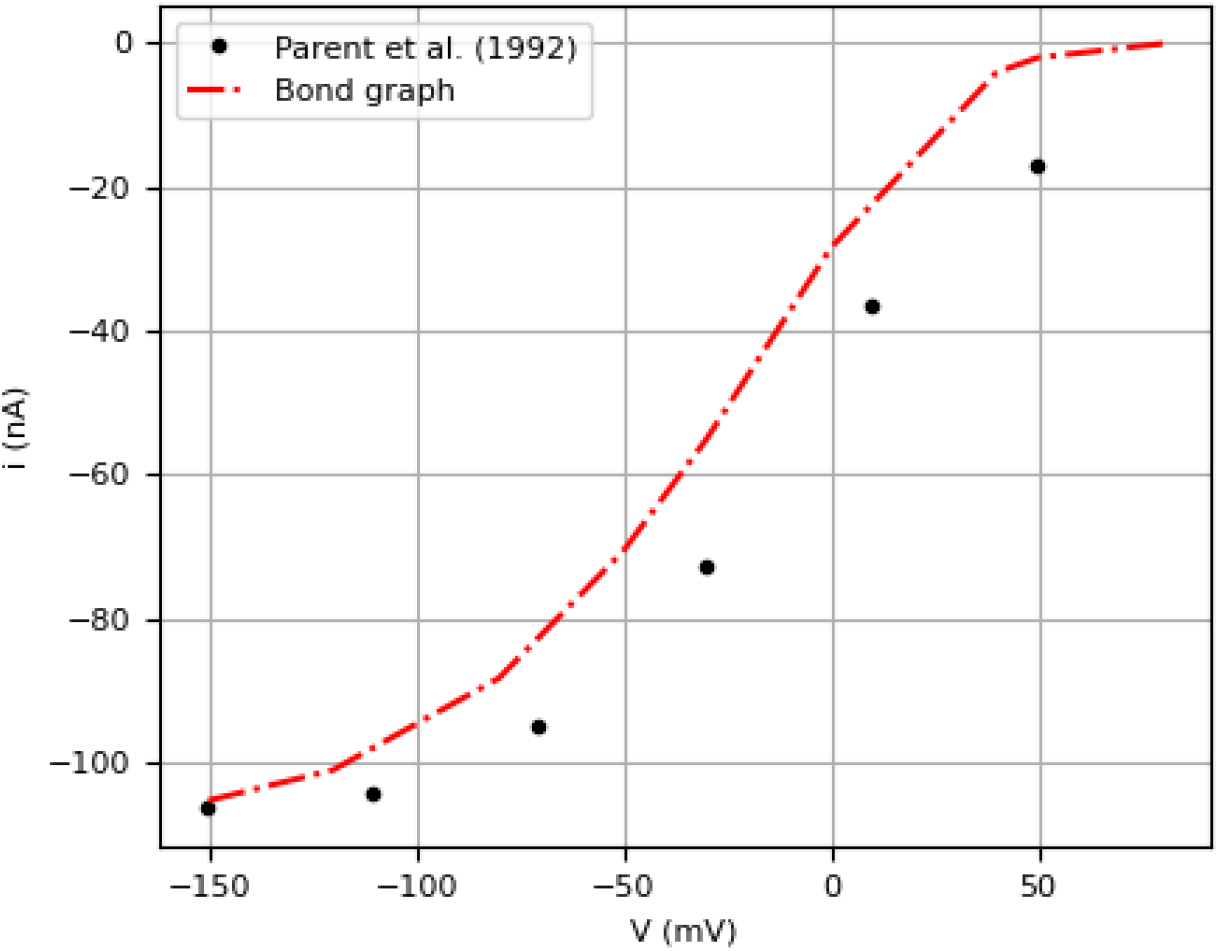
The full bond graph model compared with Figure 5 in Parent et al. [19]. The information needed to reproduce these results can be found in [18].

Note that details on the numerical implementation are given in the accompanying *Physiome* paper [18] and the code, referenced in that publication, is available on PMR.

## DISCUSSION

Bond graph models of molecular pathways have previously been developed for enzyme-catalysed reactions including glycolysis and the SLC transporter SGLT1 (but without considering the electrogenic nature of this transporter) [23], and for membrane ATPase transporters including and the cardiac sarcoplasmic/endoplasmic Ca^2+^ ATPase (SERCA) and the cardiac Na^+^/K^+^ ATPase [25]. An electrogenic version of a bond graph-based model of SGLT1 is also available [24].

In this paper we have focussed on developing a comprehensive framework for bond graph modelling of lumped parameter biological processes using the six physical units needed to represent energy transmission, storage and exchange between the mechanical, electromagnetic and chemical forms of energy (including energy dissipation to high entropy heat). We have proposed a new way of pictorially showing all components of these processes such that the equations representing conservation of mass, charge and energy, respectively, together with their constitutive laws, uniquely defined by the bond graph diagram, are easily understood by physiologists and biologists generally. We presented three examples - a coupled electromechanical actuator, a voltage-sensitive and mechano-sensitive gated ion channel, and an enzyme-catalysed reaction – to show how the bond graph framework can be used to represent all types of energy exchange.

We then developed new bond graph models for two glucose transport members of the SLC transporter family, one for facilitated diffusion (*SLC2A2*/GLUT2) and one for sodium-glucose cotransport (*SLC5A1*/SGLT1). In each case we derived the full kinetic model from the bond graph diagram and then derived a reduced steady-state model under the assumption that the binding and unbinding reactions are much faster than the reactions representing enzyme transition between inward- and outward-facing states of the transporter protein and that, because the substrates are present in much greater amounts than the transporter protein, the enzyme cycling rate can be assumed to be constant. For the second transporter, we must also assume that the slippage mechanism (protein state transition and energy dissipation with no useful transport), is not significant. The steady-state analytic models provide thermodynamically consistent generalisations of Michaelis-Menten models.

The parameters of the kinetic model and the steady-state reduced model were fitted to experimental data from the literature for each of the two types of SLC glucose transporter.

The SLC superfamily of transporter-encoding genes currently includes over 400 members with 62 families, each dealing with one specific type of transported molecule [1]. The SLC2 family, for example, deals with facilitated transport of glucose (and in one case urate), while the SLC5 family deals with sodium-assisted transport of glucose, myo-inositol, iodide, choline, lactate, or mannose. Figure 13 shows a range of these SCL transport proteins, grouped by the number of different molecules being transported and the direction of transport.

**Figure 13.**
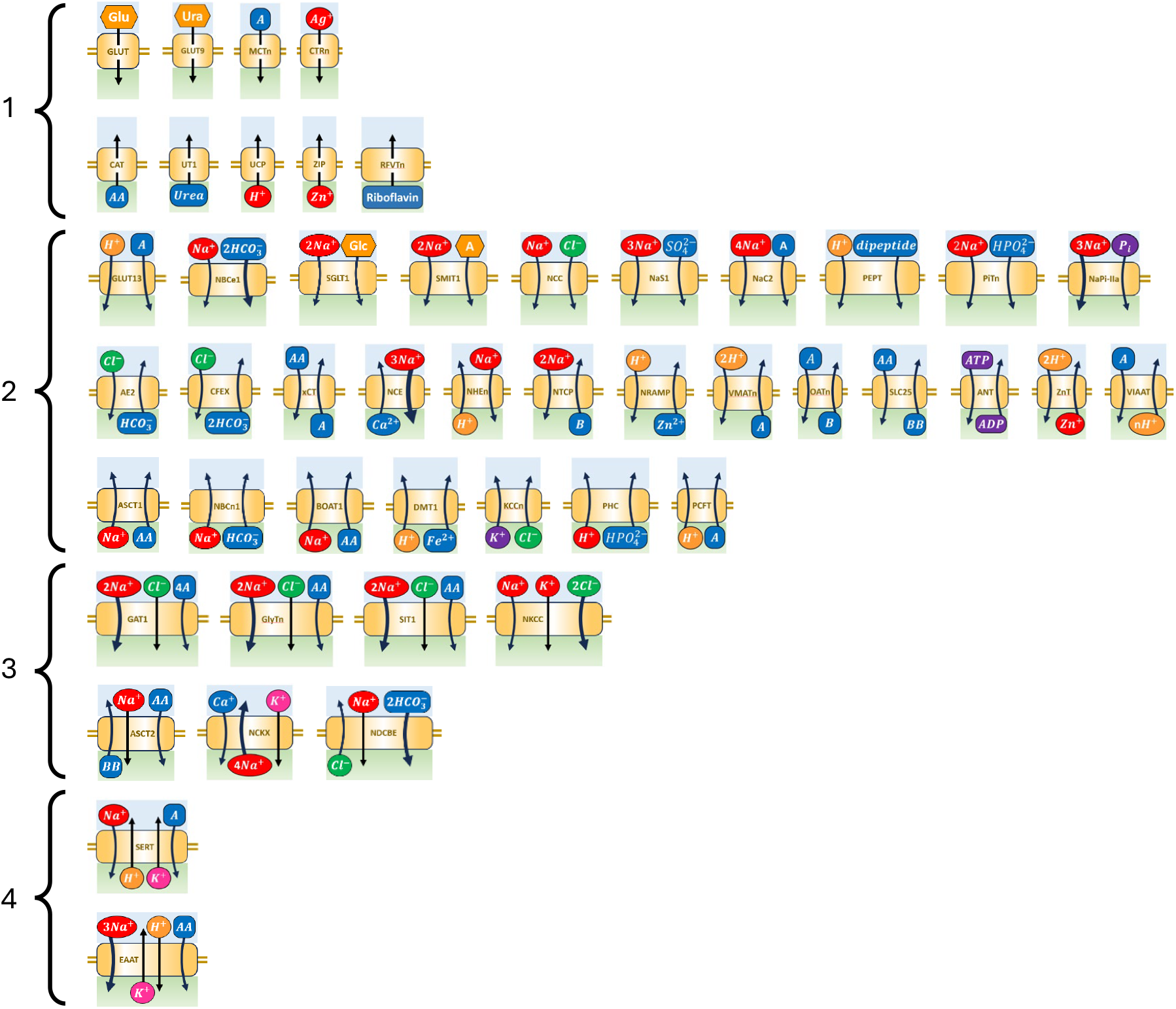
Members of the SLC transporter superfamily, grouped into four classes based on the number of solute ligands being transported. Rows within each of these four classes show examples of inwardly-directed, outwardly-directed, and two-way transport.

The extent to which the equations derived above for *SLC2A2* and *SLC5A1* can be generalised to cover a wider group of family members depends on two key factors: the sequence of binding and unbinding and, for the transport of charged molecules, the movement of charge within the membrane.

The ability to reduce the kinetic model to a steady-state relationship between flux and solute amounts is dependent on the assumption of rapid binding and unbinding, the assumption that the enzyme cycles at a constant rate (a consequence of the Briggs-Haldane assumption of relatively low expression levels for the membrane proteins compared with the availability of solute ligands), and the assumption that slippage mechanisms are not important. Each of these assumptions needs to be validated for a specific transport protein, as we have done in the examples presented here by comparing the output of the reduced model with the output of the full kinetic model (Figures 8 and 10). Note that in the approach described here we are assuming that each transporter, belonging to a template appropriate to one of the four families illustrated in Figure 13, can be fitted to flux measurements under controlled perturbations of the molar quantities of ligand on either side of the membrane. Such measurements have yet to carried out on many of these transporters. Ideally these reaction parameters would be predicted by the three-dimensional structure of the proteins (and a knowledge of the composition of the membrane sugars).

The SLC transporters in a particular cell influence one another if they transport a common ligand (such as sodium) and for electrogenic proteins there is also crosstalk via changes in the electrical potential of the membrane. We will examine composite bond graph models involving more than one transporter in future publications. See [26, 27] for modular bond graph approaches to systems biology.

## CONCLUSION

Physiological processes almost always involve energy exchange between mechanics, electromagnetics and biochemistry. We demonstrate how energy-based bond graphs can capture the processes and generate models obeying the three conservation laws of physics, particularly where these models involve the exchange of energy between the three different physical energy storage mechanisms. We developed a number of generic bond graph templates for the SLC superfamily, and fitted parameters for *SLC2A2* and *SLC5A1* to experimental data. This framework can be extended to encompass any lumped parameter physiological processes and to higher dimensional systems via port-Hamiltonians [28]. The bond graph representation of biological processes will serve as the foundation upon which high-level physiological systems will be built.

## AUTHOR CONTRIBUTIONS

P.H. and D.N. conceptualized and designed the research. W.A. implemented the model and analysed the data. All authors contributed to the manuscript and approved the submitted version.

## ACKNOWLEDGEMENTS

This project is supported by a grant from the NZ Government’s MBIE Catalyst Fund (the 12 Labours project) and a grant from the European Commission under the Horizon Europe scheme (the VITAL project). We would also like to acknowledge the huge contribution made to the use of bond graphs in physiological modelling by Edmund Crampin (who tragically died two years ago), Peter Gawthrop and Michael Pan, at the University of Melbourne.

## References

[1] M. A. Hediger, B. Clémençon, R. E. Burrier and E. A. Bruford, “The ABCs of membrane transporters in health and disease (SLC series): introduction,” Molecular aspects of medicine, vol. 34, p. 95–107, 2013.

[2] H. M. Paynter, “Analysis and design of engineering systems,” MIT press, 1961.

[3] G. F. Oster, A. S. Perelson and A. Katchalsky, “Network Thermodynamics,” Nature, vol. 234, p. 393–399, 1971.

[4] G. F. Oster, A. S. Perelson and A. Katchalsky, “Network thermodynamics: dynamic modelling of biophysical systems,” Quarterly reviews of Biophysics, vol. 6, p. 1–134, 1973.

[5] P. J. Gawthrop and E. J. Crampin, “Energy-based analysis of biochemical cycles using bond graphs,” Proceedings of the Royal Society A: Mathematical, Physical and Engineering Sciences, vol. 470, p. 20140459, 2014.

[6] P. J. Gawthrop, J. Cursons and E. J. Crampin, “Hierarchical bond graph modelling of biochemical networks,” Proceedings of the Royal Society A: Mathematical, Physical and Engineering Sciences, vol. 471, p. 20150642, 2015.

[7] H. M. Sauro, Enzyme kinetics for systems biology, Future Skill Software, 2011.

[8] J. P. Keener and J. Sneyd, Mathematical physiology, Springer New York, NY, USA, 2009.

[9] W. F. Boron and E. L. Boulpaep, Medical physiology E-book, Elsevier Health Sciences, 2016.

[10] M. Mueckler and B. Thorens, “The SLC2 (GLUT) family of membrane transporters,” Molecular aspects of medicine, vol. 34, p. 121–138, 2013.

[11] A. Carruthers, J. DeZutter, A. Ganguly and S. U. Devaskar, “Will the original glucose transporter isoform please stand up!,” American Journal of Physiology-Endocrinology and Metabolism, vol. 297, p. E836–E848, 2009.

[12] F. R. Gorga and G. E. Lienhard, “Equilibriums and kinetics of ligand binding to the human erythrocyte glucose transporter. Evidence for an alternating conformation model for transport,” Biochemistry, vol. 20, p. 5108–5113, 1981.

[13] A. G. Lowe and A. R. Walmsley, “The kinetics of glucose transport in human red blood cells,” Biochimica et Biophysica Acta (BBA)-Biomembranes, vol. 857, p. 146–154, 1986.

[14] S. Cao, Y. Chen, Y. Ren, Y. Feng and S. Long, “GLUT1 biological function and inhibition: research advances,” Future Medicinal Chemistry, vol. 13, p. 1227–1243, 2021.

[15] R. J. Naftalin, “Alternating carrier models of asymmetric glucose transport violate the energy conservation laws,” Biophysical journal, vol. 95, p. 4300–4314, 2008.

[16] J.-Y. Lapointe, L. J. Sasseville and J.-P. Longpré, “Alternating carrier models and the energy conservation laws,” Biophysical journal, vol. 97, p. 2648–2650, 2009.

[17] M. Pan, “A bond graph approach to integrative biophysical modelling.,” 2019.

[18] W. Ai, P. J. Hunter and D. P. Nickerson, “Energy-based bond graph models of glucose transport with SLC transporters,” Figshare, pp. 1–19. doi: 10.17608/k6.auckland.26073757.v3, 2024.

[19] L. Parent, S. Supplisson, D. D. F. Loo and E. M. Wright, “Electrogenic properties of the cloned Na+/glucose cotransporter: II. A transport model under nonrapid equilibrium conditions,” The Journal of membrane biology, vol. 125, p. 63–79, 1992.

[20] E. M. Wright, D. D. F. Loo and B. A. Hirayama, “Biology of human sodium glucose transporters,” Physiological reviews, vol. 91, p. 733–794, 2011.

[21] G. Gyimesi, J. Pujol-Giménez, Y. Kanai and M. A. Hediger, “Sodium-coupled glucose transport, the SLC5 family, and therapeutically relevant inhibitors: from molecular discovery to clinical application,” P?ügers Archiv-European Journal of Physiology, vol. 472, p. 1177–1206, 2020.

[22] X.-Z. Chen, M. J. Coady, F. Jackson, A. Berteloot and J.-Y. Lapointe, “Thermodynamic determination of the Na+: glucose coupling ratio for the human SGLT1 cotransporter,” Biophysical Journal, vol. 69, p. 2405–2414, 1995.

[23] P. J. Gawthrop and E. J. Crampin, “Energy-based analysis of biomolecular pathways,” Proceedings of the Royal Society A: Mathematical, Physical and Engineering Sciences, vol. 473, p. 20160825, 2017.

[24] P. J. Gawthrop and M. Pan, “Network thermodynamical modeling of bioelectrical systems: a bond graph approach,” Bioelectricity, vol. 3, p. 3–13, 2021.

[25] M. Pan, P. J. Gawthrop, K. Tran, J. Cursons and E. J. Crampin, “A thermodynamic framework for modelling membrane transporters,” Biophysical Journal, vol. 116, p. 420a, 2019.

[26] M. Pan, P. J. Gawthrop, J. Cursons and E. J. Crampin, “Modular assembly of dynamic models in systems biology,” PLoS computational biology, vol. 17, p. e1009513, 2021.

[27] P. J. Gawthrop and E. J. Crampin, “Modular bond-graph modelling and analysis of biomolecular systems,” IET Systems Biology, vol. 10, p. 187–201, 2016.

[28] F. J. Argus, C. P. Bradley and P. J. Hunter, “Theory and implementation of coupled port-Hamiltonian continuum and lumped parameter models,” Journal of Elasticity, vol. 145, p. 339–382, 2021.

